# CD20 is a mammalian odorant receptor expressed in a subset of olfactory sensory neurons that mediates innate avoidance of predators

**DOI:** 10.1101/2023.08.08.552498

**Authors:** Hao-Ching Jiang, Sung Jin Park, I-Hao Wang, Daniel M. Bear, Alexandra Nowlan, Paul L. Greer

## Abstract

The mammalian olfactory system detects and discriminates between millions of odorants to elicit appropriate behavioral responses. While much has been learned about how olfactory sensory neurons detect odorants and signal their presence, how specific innate, unlearned behaviors are initiated in response to ethologically relevant odors remains poorly understood. Here, we show that the 4-transmembrane protein CD20, also known as MS4A1, is expressed in a previously uncharacterized subpopulation of olfactory sensory neurons in the main olfactory epithelium of the murine nasal cavity and functions as a mammalian odorant receptor that recognizes compounds produced by mouse predators. While wild-type mice avoid these predator odorants, mice genetically deleted of CD20 do not appropriately respond. Together, this work reveals a novel CD20-mediated odor-sensing mechanism in the mammalian olfactory system that triggers innate behaviors critical for organismal survival.

## Introduction

To survive, animals must accurately detect, correctly interpret, and appropriately respond to sensory stimuli in their environment. For most non-primate mammals, the richest source of this information is the immense variety of small molecules present in their external surroundings, which may signify the presence of predators, food, or mates (Brennan and Zufall, 2006). These chemicals are primarily detected by odorant receptors (ORs) expressed at the sensory endings of peripheral olfactory sensory neurons (OSNs), which are coupled to the higher brain circuits tasked with mediating odor perception and initiating olfactory-driven behavior (Bargmann, 2006; Buck and Axel, 1991; Leinwand and Chalasani, 2011; Su et al., 2009). However, how mammals detect and process different classes of olfactory stimuli to initiate distinct behaviors is still not well understood (*i.e*., how does a mouse know to avoid a cat but to actively seek out a piece of cheese?).

One emerging hypothesis is that distinct subpopulations of OSNs might be responsible for different behaviors. The olfactory system can be subdivided into multiple anatomically and molecularly distinct subpopulations of OSNs. In the mouse there are at least nine distinct olfactory subsystems, each of which is made up of unique, and non-overlapping, collections of OSNs (Bargmann, 1997; Hu et al., 2007; Liberles and Buck, 2006; Lin et al., 2007; Liu et al., 2009; Omura and Mombaerts, 2014, 2015; Rivière et al., 2009; Shinoda et al., 1989). A handful of these olfactory subsystems have been extensively studied, which has led to significant insight into their role in olfactory perception and odor-driven behaviors. Particularly important for elucidating the role of these olfactory subsystems has been the identification of the ORs that they express. Each subsystem expresses different types of ORs, which enable them to detect subsets of chemical space and mediate specific behaviors. For instance, the largest subdivision in the mouse, the main olfactory system, owing to its immense receptor repertoire of approximately 1000 distinct ORs (Buck and Axel, 1991), is able to detect essentially all volatile odorants and therefore plays a key role in odor discrimination and odorant-dependent learning (Kajiya et al., 2001; Sanchez-Andrade and Kendrick, 2009; Touhara, 2002). Smaller subsystems, such as the vomeronasal subsystem and the trace amine-associated receptor (TAAR) subsystem express much smaller receptor repertoires that are more narrowly tuned to recognize specific classes of behaviorally relevant odorants (Dulac and Axel, 1995; Liberles and Buck, 2006) and may therefore have more specialized roles in identifying odors of innate significance and initiating specific patterns of unlearned behaviors critical for survival (Del Punta et al., 2002; Dewan et al., 2013; Liberles and Buck, 2006).

Nonetheless, despite progress in elucidating the function of a few of these olfactory subsystems, the specific roles of others remain poorly understood. One of the least understood is the olfactory necklace subsystem, which seems to mediate seemingly opposing behaviors for both feeding and innate avoidance of noxious stimuli (Hu et al., 2007; Munger et al., 2010). Perhaps the biggest hurdle toward understanding the role of the necklace system in odor-driven behavior is that until recently it was unclear how it detects odorants. We identified the membrane spanning 4A (MS4A) family of proteins as a novel set of ORs in mammalian necklace OSNs (Greer et al., 2016). Heterologous expression of individual MS4A proteins in HEK293 cells conferred the ability to respond to specific chemical compounds. Moreover, the *in vitro* odor receptive fields of MS4A proteins matched those of the necklace OSNs in which MS4A proteins were expressed (Greer et al., 2016). Nonetheless, an absence of mouse lines in which *Ms4a* gene expression was genetically manipulated meant that the role of MS4A proteins in necklace olfactory function was only examined in *in vitro* and *ex vivo* experiments, therefore preventing a rigorous assessment of whether MS4A proteins participate in odor detection *in vivo*. Indeed, because MS4A proteins do not resemble any previously described odorant receptors – they are four-transmembrane spanning proteins rather than seven-transmembrane GPCRs, there remains some skepticism about whether MS4A proteins function as ORs *in vivo* (Zimmerman and Munger, 2021). Here, we use newly generated *Ms4a* knockout mice to show that MS4A proteins function as *bona fide* ORs *in vivo.* Moreover, we show that the MS4A family member MS4A1, (better known as CD20, a protein previously identified as a co-receptor for the B cell receptor in lymphocytes), is not expressed in the necklace, but is instead expressed in a novel subset of OSNs outside the necklace. Within this subpopulation of OSNs, MS4A1 senses predator odorants leading to innate, unlearned avoidance behaviors.

## Results

### MS4A proteins function as chemoreceptors in necklace OSNs *in vivo* and mediate specific odor-driven avoidance behaviors

Our previous work suggested that *Ms4a* genes encode a new family of non-GPCR mammalian ORs (Greer et al., 2016). However, a lack of genetically modified mice in which *Ms4a* expression could be manipulated precluded a definitive determination of whether *Ms4a* genes do in fact encode *bona fide* ORs that function *in vivo*. Addressing this issue is particularly important given the unusual structure and expression pattern of MS4A proteins in the mammalian olfactory system (Greer et al., 2016). To circumvent potential issues of redundancy between *Ms4a* family members, we took advantage of a mouse in which CRISPR/Cas9 technology was deployed to delete all 17 murine *Ms4a* genes (hereafter, referred to as *Ms4a* cluster knockout mice) (Figures 1A, S1A, and S1B). *Ms4a* cluster knockout mice are viable, fertile, produced at Mendelian frequency, and are overtly indistinguishable from their wild-type littermates, enabling us to assess olfactory performance in these *Ms4a*-deficient animals (Figure S1C).

**Figure 1.**
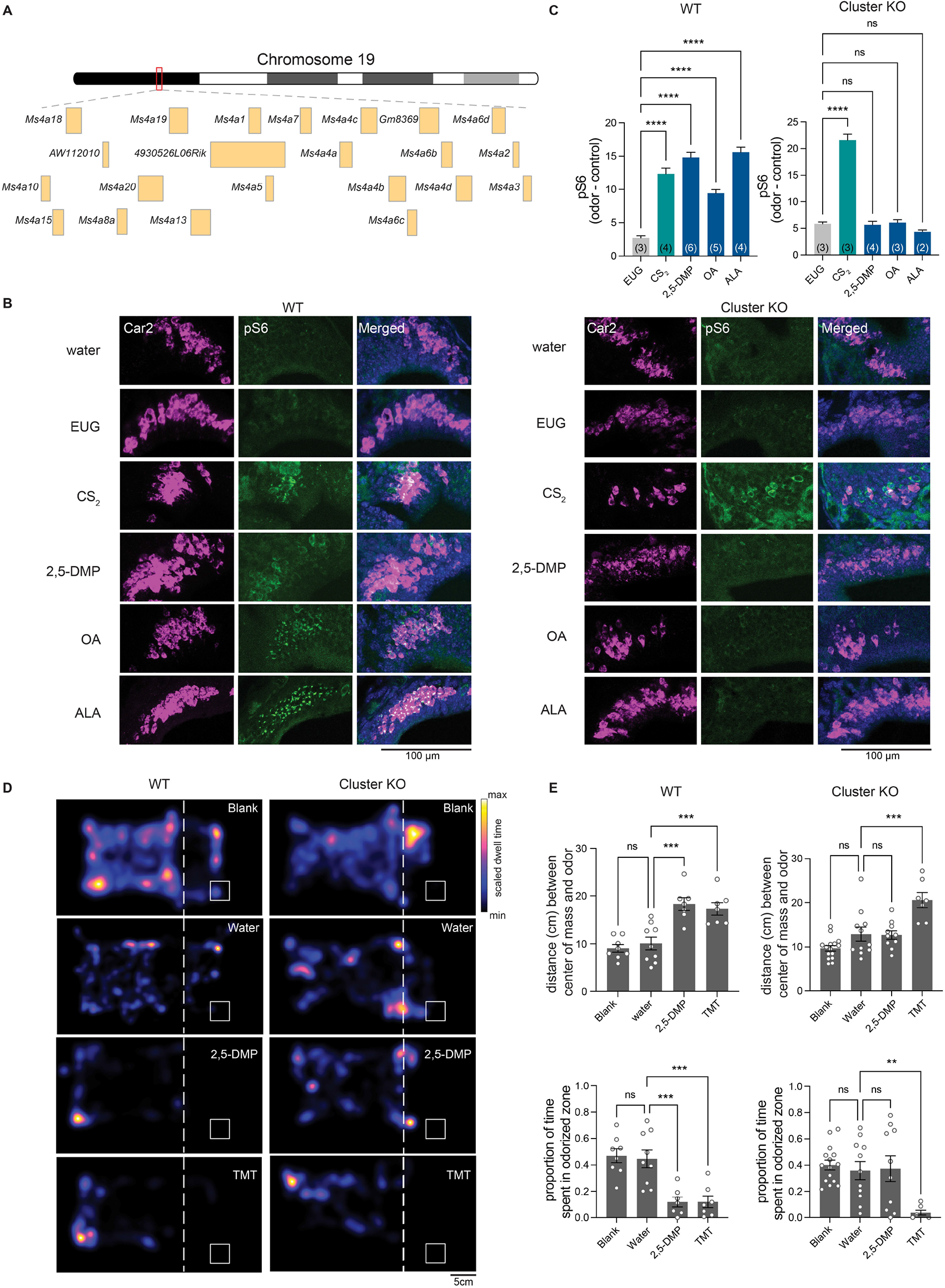
Deletion of all members of the MS4A family prevents *in vivo* detection of MS4A ligands by necklace OSNs and the innate avoidance behaviors trigged by these ligands. (A) Schematic representation of the genomic organization of the 19 Ms4a family members in tandem array on chromosome 19. (B) Example images of the cul-de-sac regions (where necklace cells reside) of the main olfactory epithelia of mice exposed to the indicated odorant, co-labeled for the necklace cell marker Car2 (purple) and the neuronal activity marker phospho-S6 (pSerine240/244) (green) from wild-type (left panels) or Ms4a cluster knockout animals (right panels). EUG; eugenol, CS_2_; carbon disulfide, 2,5-DMP; 2,5-dimethylpyrazine, OA; oleic acid, ALA; alpha-linolenic acid. (C) Quantification of the proportion of pS6+ necklace cells in odor-exposed wild-type mice (left panel) or Ms4a cluster knockout mice (right panel). Data are presented as mean ± SEM, n ≥ three independent experiments, ****p < 0.0001, Dunnett’s test following one-way ANOVA compared to null exposure. TMT; 2,3,5-trimethyl-3-thiazoline. (D) Heat maps of the occupancy of wild-type (left panels) or cluster knockout mice (right panels) in the odor avoidance chamber in response to the indicated odorants. Small square represents location of odorant, and dashed line demarcates the odor avoidance zone from the rest of the chamber. Scale bar, 5 cm. (E) Quantification of odor avoidance behavior. The distance between the average center of mass of the mouse and the location of odorant (top panels) and the proportion of time spent in the odorized zone (bottom panels) were determined for wild-type mice (left) and Ms4a cluster knockout mice (right). Each circle represents an individual mouse. Data are presented as mean ± SEM, n ≥ five independent experiments, **p < 0.01, ***p < 0.001, unpaired Welch t-test or post-hoc Dunnett’s test following one-way ANOVA or a paired t-test compared to water exposure.

To begin to test the role of *Ms4a* genes in olfactory function, we initially exposed freely behaving *Ms4a*-deficient mice or their wild-type littermates to 2,5-dimethylpyrazine (2,5-DMP), oleic acid (OA), or alpha-linolenic acid (ALA), previously described *in vitro* ligands of MS4A6C, MS4A6D, and MS4A4B, respectively, and measured activation of necklace OSNs, which express MS4A proteins (Greer et al., 2016), by detecting S6 phosphorylation (pS6), a well-established marker of OSN activation (Jiang et al., 2015). Deletion of *Ms4a* genes eliminated necklace OSN pS6 responses to each of the MS4A-triggering compounds (Figures 1B and 1C). By contrast, *Ms4a* cluster knockout mice necklace cells responded without impairment to carbon disulfide (CS_2_), which is detected by necklace cells through the actions of the receptor guanylate cyclase, GCD, in an MS4A-independent manner (Figures 1B and 1C). Thus, *Ms4a* deletion did not disrupt the health of necklace cells or their capacity to respond to odors in general, but instead specifically prevented their detection of MS4A ligands suggesting that MS4A proteins function as ORs in necklace OSNs *in vivo*.

Next, we wanted to examine what role, if any, MS4A odorant receptors play in odor-driven behavior. The necklace olfactory system in which *Ms4a* genes are expressed has been implicated in two distinct MS4A-independent olfactory behaviors—the innate avoidance of CO_2_ at concentrations above those naturally found within the atmosphere (Hu et al., 2007), and the social transmission of food avoidance triggered by CS_2_ (Munger et al., 2010) or the urinary peptides, guanylin and uroguanylin (Leinders-Zufall et al., 2007). Each of these behaviors is thought to be mediated through the receptor guanylate cyclase, GCD (Leinders-Zufall et al., 2007; Munger et al., 2010; Sun et al., 2009). As MS4A receptors are also expressed in necklace OSNs (Greer et al., 2016), we sought to determine whether MS4A receptors contribute to similar types of innate odor-driven behaviors.

We initially focused our efforts on determining whether MS4A receptors mediate innate, odor-driven avoidance responses as these behaviors are robust, reproducible, do not require prior training, and are easily quantifiable. To begin to determine whether MS4As mediate innate avoidance behaviors, we first tested whether any previously identified MS4A ligands induce avoidance in wild-type mice in an unlearned manner. 2,3-dimethylpyrazine (2,3-DMP), 2,5-DMP, OA, linolenic acid (LA), ALA, and arachidonic acid (AA) all activate MS4A-expressing cells *in vitro* and necklace OSNs *in vivo* (Greer et al., 2016). 2,3-DMP and 2,5-DMP are found in the urine of wolves and ferrets, natural mouse predators, respectively, and prior work has suggested that these compounds are ethologically relevant odors for mice (Apfelbach et al., 2015; Brechbühl et al., 2013; Osada et al., 2014; Zhang et al., 2005). By contrast, OA, LA, ALA, and AA are long chain fatty acids found in natural food sources of mice (Abedi and Sahari, 2014; Cordova et al., 2012). To determine whether any of these MS4A ligands trigger innate avoidance responses, we compared the aversive behavior of *Ms4a* cluster knockout and wild-type mice in response to these compounds and to 2,3,5-trimethyl-3-thiazoline (TMT), a component of fox feces (Fendt et al., 2005) that is not an MS4A ligand. The MS4A ligands 2,3-DMP and 2,5-DMP as well as the previously described aversive odorant, TMT, induced innate, unlearned avoidance responses in wild-type mice (Figures 1D, 1E and S1D). Although wild-type mice robustly avoided DMP, *Ms4a* cluster knockout mice were oblivious to DMP and behaved as though no odor was present (Figures 1D and 1E). This effect of *Ms4a* deletion on mouse avoidance behavior was specific to DMP; *Ms4a* cluster knockouts exhibited similar aversive behaviors as their wild-type littermates in response to other ethologically relevant aversive odors that are not MS4A ligands, such as TMT (Figures 1D and 1E). In addition, deletion of *Ms4as* did not affect other non-odor mediated avoidance behaviors such as the amount of time spent in open arms in an elevated plus maze assay (Figure S1E). Taken together, these results indicate that *Ms4a* genes encode ORs that mediate specific odor-driven avoidance responses in mammals.

### Ms4a6c detects DMP in necklace OSNs *in vivo* but does not fully mediate avoidance behaviors to DMP

We next sought to determine which of the 17 *Ms4a* family members were responsible for mediating the unlearned avoidance mice exhibit in response to DMP. Because DMP is sensed by MS4A6C *in vitro* (Greer et al., 2016), we initially focused on this MS4A family member and utilized a mouse line in which the *Ms4a6c* gene was specifically deleted (Figures S2A and S2B). Like *Ms4a* cluster knockout mice, *Ms4a6c* knockout mice were viable, fertile, and overtly indistinguishable from their wild-type littermates (Figures S2C and S2D). Consistent with previous work demonstrating that MS4A6C detects DMP *in vitro*, *Ms4a6c* knockout necklace neurons did not respond to 2,3-DMP or 2,5-DMP as assessed by pS6 staining (Figures 2A and 2B). By contrast, necklace neurons of *Ms4a6c* knockout mice still robustly responded to OA, the *in vitro* ligand of the closely related MS4A family member, MS4A6D (Greer et al., 2016) and to CS_2_, a GCD ligand (Munger et al., 2010), indicating that *Ms4a6c* deletion specifically impairs the ability of necklace cells to detect 2,3-DMP and 2,5-DMP and does not generally disrupt their ability to sense non-MS4A6C odorants (Figures 2A and 2B).

**Figure 2.**
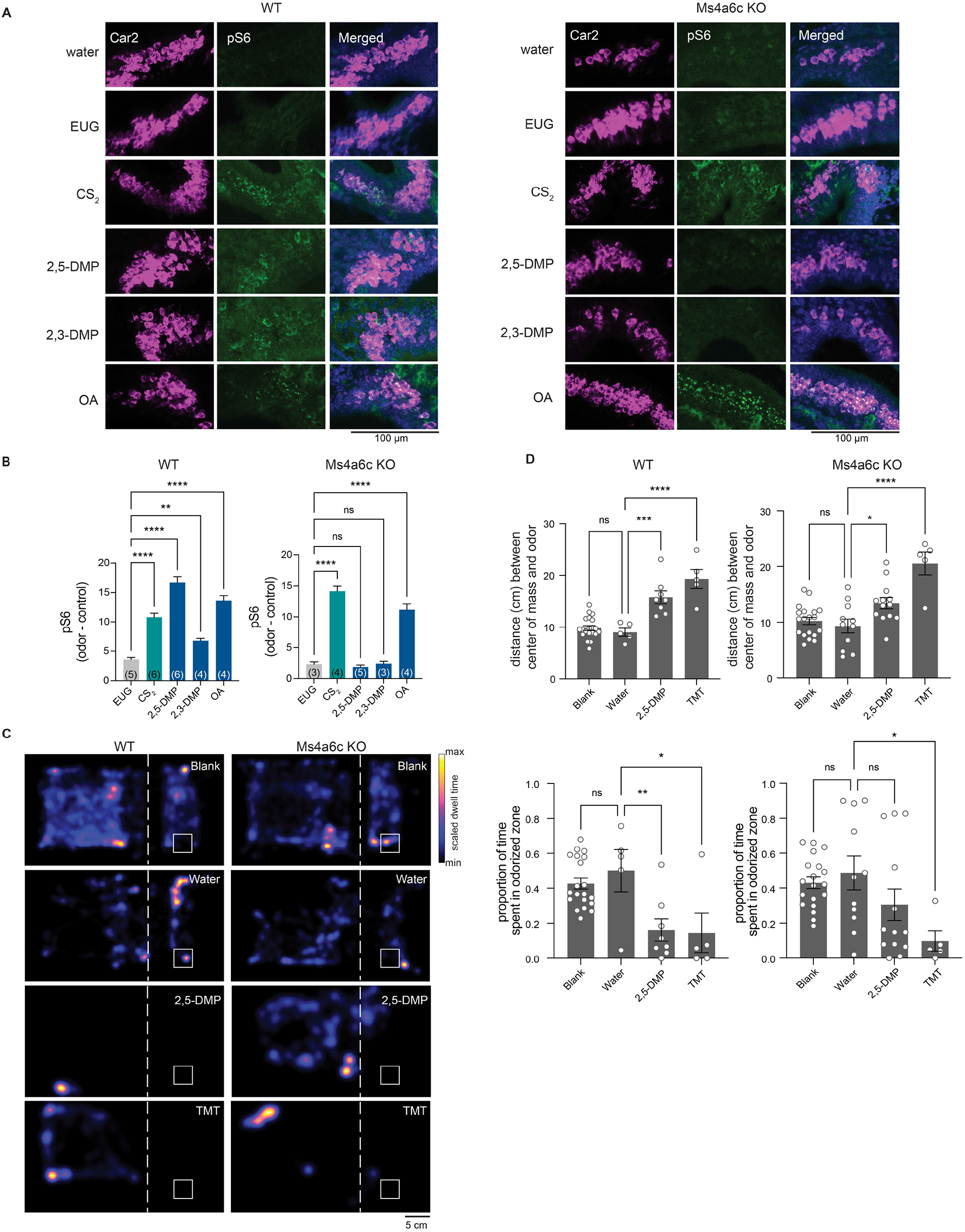
The knockout of *Ms4a6c* impairs the ability of necklace OSNs to detect DMP, a predator-derived compound, but does not significantly affect DMP-mediated innate avoidance behavior. (A) Example images of the cul-de-sac regions of the main olfactory epithelia of mice exposed to the indicated odorant, co-labeled for the necklace cell marker, Car2 (purple), and the neuronal activity marker, phospho-S6 (pSerine240/244) (green), from wild-type (left panels) or *Ms4a6c* knockout animals (right panels). EUG; eugenol, CS_2_; carbon disulfide, 2,3-DMP; 2,3-dimethylpyrazine, 2,5-DMP; 2,5-dimethylpyrazine, OA; oleic acid. (B) Quantification of the proportion of pS6+ necklace cells in odor-exposed wild-type mice (left panel) or *Ms4a6c* knockout mice (right panel). Data are presented as mean ± SEM, n ≥ three independent experiments, **p < 0.01, ****p < 0.0001, Dunnett’s test following one-way ANOVA to null exposure. (C) Heat maps of the occupancy of wild-type (left panels) or *Ms4a6c* knockout mice (right panels) in the odor avoidance chamber in response to the indicated odorants. Small square represents location of odorant, and dashed line demarcates the odor avoidance zone from the rest of the chamber. Scale bar, 5 cm. TMT; 2,3,5-trimethyl-3-thiazoline. (D) Quantification of odor avoidance behavior. The distance between the average center of mass of the mouse and the location of odorant (top panels) and the proportion of time spent in the odorized zone (bottom panels) were determined for wild-type mice (left) and *Ms4a6c* knockout mice (right). Each circle represents an individual mouse. Data are presented as mean ± SEM, n ≥ five independent experiments, *p < 0.05, **p < 0.01, ***p < 0.001, ****p < 0.0001, unpaired Welch t-test or post-hoc Dunnett’s test following one-way ANOVA or a paired t-test compared to water exposure.

To determine whether the failure of necklace cells to detect MS4A6C ligands altered avoidance of these odorants, we assessed innate avoidance responses to 2,5-DMP by *Ms4a6c*-deficient mice. Surprisingly, although *Ms4a6c* knockout mice avoided 2,5-DMP somewhat less than wild-type mice, they avoided 2,5-DMP significantly more than *Ms4a* cluster knockout mice (Figures 2C, 2D, and 3E). This result suggests that at least one additional *Ms4a* family member may mediate innate avoidance of DMP.

### CD20 responds to DMP and mediates DMP-driven innate avoidance behaviors

To identify additional MS4A receptor(s) that sense DMP, we assessed the ability of all 17 murine MS4A family members to detect 2,5-DMP by detecting DMP-induced calcium responses to odorants in HEK293 cells co-expressing individual *Ms4a* genes with the genetically encoded, fluorescent calcium indicator, GCaMP6s. HEK293 cells do not express endogenous MS4A proteins (Greer et al., 2016), but exogenously expressed MS4A proteins are efficiently trafficked to the plasma membrane within these cells (Figure S3A). HEK293 cells expressing either MS4A6C or MS4A1, but none of the other MS4A family proteins, responded to 2,5-DMP to generate a transient calcium signal (Figures 3A and 3B), suggesting that MS4A1 is the other MS4A family member mediating the mouse’s innate avoidance response to 2,5-DMP. To test this hypothesis, we assessed the ability of *Ms4a1* knockout mice to avoid 2,5-DMP. *Ms4a1* knockout mice acted like *Ms4a* cluster knockout mice – exhibiting no avoidance responses to this predator-derived compound (Figures 3C-E). The failure of *Ms4a1*-deficient mice to respond to 2,5-DMP was specific to this odor since *Ms4a1* knockout mice avoided other aversive odorants such as TMT to the same extent as wild type mice (Figures 3C and 3D). Moreover, *Ms4a1* knockout mice were overtly indistinguishable from wild type mice in other ways – they exhibited similar locomotive behaviors and behaved similarly to wild-type mice in assays of anxiety (such as the elevated plus maze) (Figures S3B and S3C), strongly suggesting that the failure to respond to 2,5-DMP was a specific defect in this particular odor-driven behavior and not a sign of more general nervous system dysfunction.

**Figure 3.**
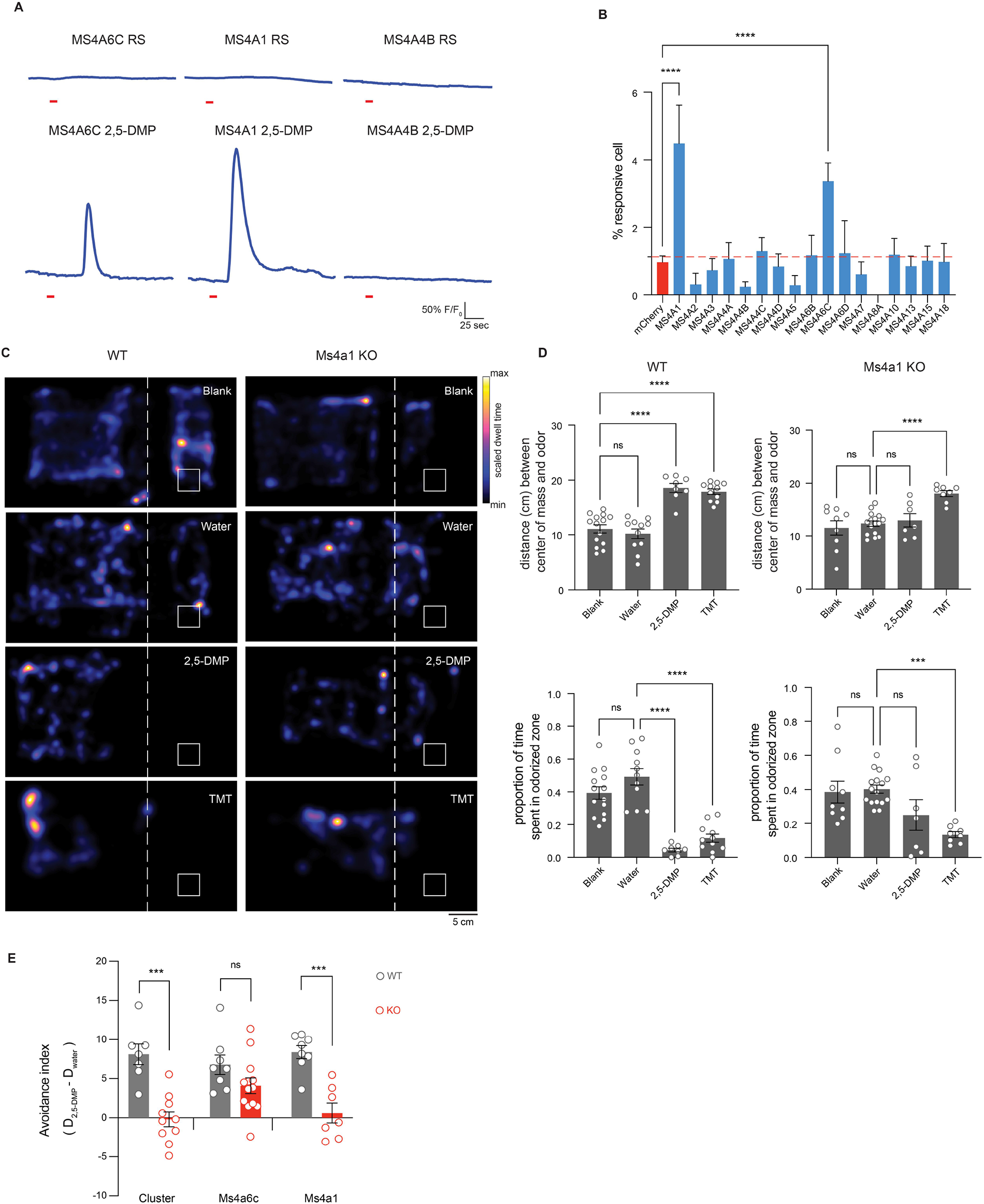
MS4A1/CD20 facilitates the detection of DMP, and deletion of *Ms4a1* eliminates innate avoidance of DMP. (A) GCaMP6s fluorescence in response to the indicated chemicals in HEK293 cells expressing the indicated MS4A protein (odor delivery indicated by red bars). RS; Ringer’s Solution, 2,5-DMP; 2,5-dimethylpyrazine. (B) Quantification of responses of expressed MS4A protein to 2,5-DMP as in (A). n ≥ 3 independent experiments. Dashed red line indicates mean plus 2.5 standard deviations above the responses of HEK293 cells expressing mCherry alone in response to 2,5-DMP. **** p < 0.0001, Dunnett’s test following one-way ANOVA compared to mCherry alone. (C) Heat maps of the occupancy of wild-type (left panels) or *Ms4a1* knockout mice (right panels) in the odor avoidance chamber in response to the indicated odorants. Small square represents location of odorant, and dashed line demarcates the odor avoidance zone from the rest of the chamber. Scale bar, 5 cm. TMT; 2,3,5-trimethyl-3-thiazoline. (D) Quantification of odor avoidance behavior. The distance between the average center of mass of the mouse and the location of odorant (top panels) and the proportion of time spent in the odorized zone (bottom panels) were determined for wild-type mice (left) and *Ms4a1* knockout mice (right). Each circle represents an individual mouse. Data are presented as mean ± SEM, n ≥ five independent experiments, ***p < 0.001, ****p < 0.0001, unpaired Welch t-test or post-hoc Dunnett’s test following one-way ANOVA or a paired t-test compared to water exposure. (E) An avoidance index was calculated for cluster knockout mice, *Ms4a6c* knockout mice, *Ms4a1* knockout mice, and their wild-type littermate controls by subtracting the average distance in cm between the average position of a mouse from water from the average position of the mouse and 2,5-DMP. A more positive value represents a larger avoidance of DMP. The avoidance index was calculated for > five mice per genotype, and the data are presented as mean ± SEM. ***p < 0.001, using an unpaired Welch t-test compared to wild-type mice.

The observation that MS4A1 is required for a mouse to avoid the predator-derived compound 2,5-DMP was surprising since the only previously ascribed function of MS4A1 is as a co-receptor for the B-cell receptor in circulating mature lymphocytes, where it is known as CD20 (Tedder and Engel, 1994; Tedder et al., 1988). Although it seemed unlikely that lymphocytes would play a critical role in mediating this olfactory-driven behavior, we assessed the ability of Rag-1-deficent mice, which lack all mature lymphocytes (Mombaerts et al., 1992), to avoid 2,5-DMP. Rag-1 knockout mice avoided DMP to a similar extent as wild-type mice, indicating that mature lymphocyte function was not required for avoidance of 2,5-DMP and further suggesting that CD20 might act in cells outside of the immune system to mediate avoidance of this odor (Figures S3D and S3E).

### CD20 is expressed in a previously unidentified subpopulation of OSNs

To identify cells in the olfactory system in which *Ms4a1* might be expressed, we stained coronal sections of the mouse olfactory epithelium with an antibody specific for MS4A1. A relatively sparse population of MS4A1-expressing cells that did not express lymphoid markers was found, whose cell bodies reside within the epithelial layer of the main olfactory epithelium (MOE) (Figure 4A). To verify this unexpected observation, we stained coronal sections of the mouse olfactory epithelium with two additional anti-MS4A1 antibodies (raised in different species and recognizing different MS4A1 epitopes). These three anti-MS4A1 antibodies all co-labeled the same cells in the MOE (Figure 4B). These antibodies did not stain any cells in olfactory epithelial sections obtained from *Ms4a1* knockout mice, confirming their specificity (Figure 4C). Moreover, combined fluorescent *in situ* hybridization and immunohistochemistry experiments detected *Ms4a1* mRNA and MS4A1 protein in the same cells indicating that *Ms4a1* is expressed in non-lymphoid cells of the mouse olfactory system (Figure 4D).

**Figure 4.**
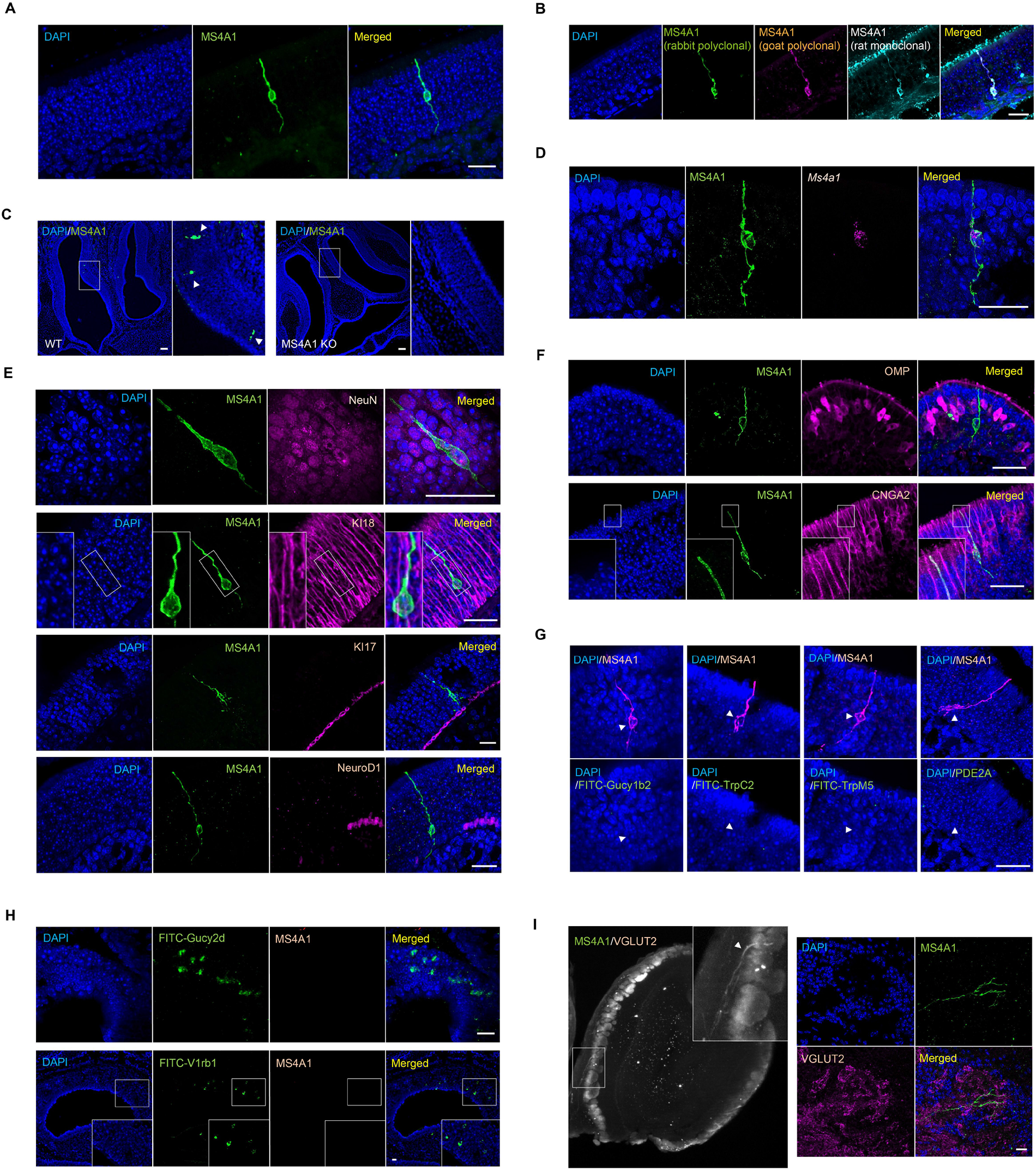
MS4A1 is expressed in a previously unidentified subpopulation of OSNs. (A) Expression of MS4A1 protein in solitary cells of the main olfactory epithelium. Scale bar, 20 μm. (B) Immunostaining performed with rabbit polyclonal (green), goat polyclonal (purple), and rat monoclonal (cyan) anti-MS4A1 antibodies recognizing different epitopes of the protein. Scale bar, 20 μm. (C) Immunostaining of MS4A1 in the main olfactory epithelia of wild-type and *Ms4a1* knockout mice. MS4A1-expressing cells are detected in sections obtained from wild-type (indicated by white arrow heads, left panels) but not *Ms4a1* knockout (right panels) mice. Scale bar, 80 μm. (D) Detection of *Ms4a1* mRNA expression (purple) in anti-MS4A1 antibody labeled cells (green) using combined single-molecule fluorescent *in situ* hybridization and immunohistochemistry. Scale bar, 20 μm. (E) Determination of whether MS4A1-expressing cells co-express NeuN (neuronal marker, top panels) KI18 (sustentacular cell marker, second panels from the top), KI17 (horizontal basal cell marker, third panels from the top), and NeuroD1 (globose basal cell marker, bottom panels). Scale bar, 20 μm. (F) Expression of CNGA2 (lower panels) but not OMP (upper panels) in MS4A1+ OSNs. Scale bar, 20 μm. (G) MS4A1-expressing cells do not express the genes Gucy1b2, Trpc2, Trpm5, and Pde2a, markers of previously described olfactory subsystems. Scale bar, 20 μm. (H) MS4A1 is undetected in necklace cells marked by their expression of Gucy2d (upper panels) and vomeronasal olfactory neurons marked by their expression of V1rb1 (lower panels). Scale bar, 20 μm. (I) iDISCO (left panel) and immunostaining (right panels) using antibodies of MS4A1 and VGLUT2, an olfactory glomerular marker, reveal that MS4A1-expressing cells coalesce their axons in the olfactory bulb.

The cell bodies of MS4A1-expressing cells resided in the same anatomic location as OSN cell bodies and extended what appeared to be sensory dendrites to the lumen of the MOE and axonal-like structures toward the olfactory bulb suggesting that MS4A1-expressing cells might be OSNs. To confirm that MS4A1-expressing cells are neurons, we co-stained for MS4A1 and the neuronal marker NeuN and found that all MS4A1-expressing cells in the olfactory epithelium also expressed NeuN (Figure 4E). Consistent with this observation, MS4A1 cells did not stain for KI18, a marker of glial support cells (Holbrook et al., 2011), KI17, a marker of horizontal basal cells (Holbrook et al., 2011), or NeuroD1, a marker of globose basal cells (Packard et al., 2011) (Figure 4E). MS4A1-expressing cells expressed CNGA2, a cyclic nucleotide gated olfactory channel found in mature OSNs (Brunet et al., 1996; Firestein, 2001), but not OMP, a marker of conventional GPCR OR-expressing OSNs (Danciger et al., 1989; Margolis, 1972) (Figure 4F). Together, these results suggest that MS4A1 is expressed in an unconventional neuronal cell type in the olfactory epithelium.

In mammals, a number of distinct subpopulations of olfactory sensory neurons have been previously identified, which are characterized by their unique anatomic and/or molecular features (Bargmann, 1997; Hu et al., 2007; Liberles and Buck, 2006; Lin et al., 2007; Liu et al., 2009; Omura and Mombaerts, 2014, 2015; Rivière et al., 2009; Shinoda et al., 1989). To determine to which, if any, of these olfactory subdivisions, the MS4A1-expressing cells we identified might belong, we performed immunohistochemical and fluorescent *in situ* hybridization analyses. MS4A1 was not expressed in Gucy1b2-expressing OSNs, TrpC2- or TrpM5-positive OSNs, Pde2a-expressing necklace OSNs (Figures 4G and 4H), or OSNs of the vomeronasal organ (Figure 4H). Nonetheless, using a combination of iDISCO tissue clearing and light sheet microscopy, we found that like other OSN populations, MS4A1-expressing neurons also extended their axons into glomeruli within the mouse olfactory bulb suggesting that MS4A1 is expressed in a previously uncharacterized population of olfactory sensory neurons in the olfactory epithelium and that like other members of the MS4A family, MS4A1 might also function as an olfactory chemoreceptor (Figure 4I).

### MS4A1 detects nitrogenous heterocyclic compounds *in vitro*

To begin to test this hypothesis, and to explore what types of odors MS4A1 might detect, we examined whether heterologously expressed MS4A1 might respond to additional extracellular chemicals to mediate a calcium influx in HEK293 cells co-expressing GCaMP6s (Figure 5A). Expression of MS4A1 did not increase the baseline rate of calcium transients in HEK293 cells (Figure S5A) but increased intracellular calcium spikes upon presentation of specific chemicals (Figures 5A and 5B). This was true for both human and mouse MS4A1 proteins (Figures 5A and S5B). MS4A1 responses were tuned to nitrogenous heterocyclic compounds, including 2,3-DMP, 2,5-DMP and 2,6-DMP, and to a lesser extent indole and quinoline (Figures 5B and S5C). However, not all nitrogenous heterocyclic compounds induced calcium transients in MS4A1-expressing cells, nor did non-nitrogenous compounds like isoamyl acetate and vanillin, indicating some ligand specificity (Figure 5B). Dose response curves revealed nanomolar and low micromolar EC50s for two specific MS4A1-ligand pairs, which is well within the range of what has been observed for other mammalian odorant receptor/ligand relationships (Figure 5C). Moreover, depletion of extracellular calcium completely eliminated all calcium transients observed in response to ligand presentation (Figure 5D). Together, these observations suggest that MS4A1 is a chemoreceptor that detects nitrogenous heterocyclic compounds.

**Figure 5.**
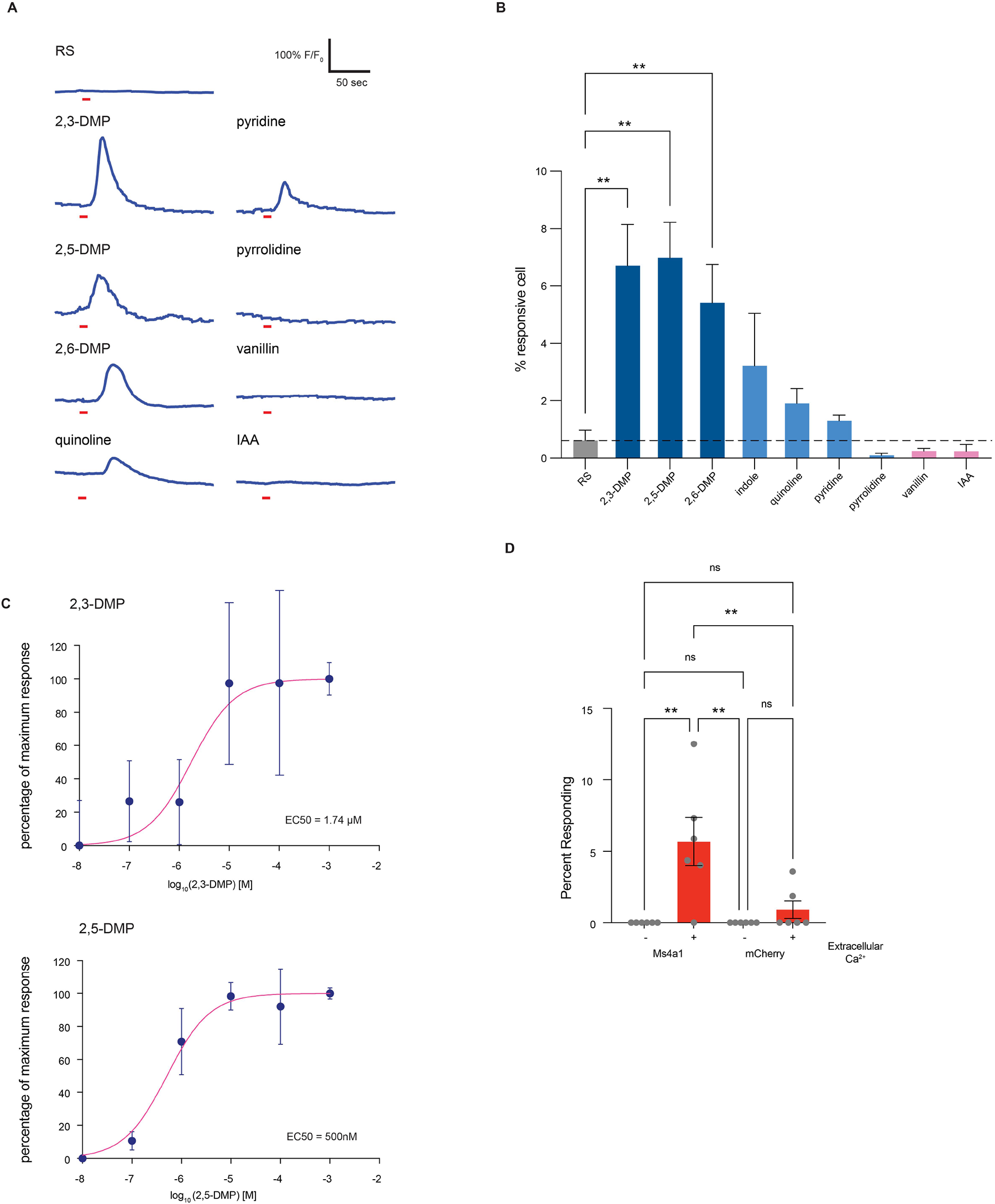
MS4A1 is a chemoreceptor that detects nitrogenous heterocyclic compounds. (A) GCaMP6s fluorescence in response to the indicated chemicals in HEK293 cells expressing MS4A1 protein (odor delivery indicated by red bars). RS; Ringer’s Solution, 2,3-DMP; 2,3-dimethylpyrazine, 2,5-DMP; 2,5-dimethylpyrazine, 2,6-DMP; 2,6-dimethylpyrazine, IAA; isoamyl alcohol. (B) Quantification of responses of cells expressing MS4A1 protein to the indicated chemicals as in (A). n ≥ three independent experiments. Dashed red line indicates mean +/-2.5 standard deviations of responses of HEK293 cells expressing mCherry alone in response to 2,5-DMP. **** p < 0.0001, Dunnett’s test following one-way ANOVA compared to mCherry alone. RS; ringer solution. (C) Dose-response curves reveal low micromolar/high nanomolar EC_50_s for MS4A1/2,3-DMP (top panel) and MS4A1/2,5-DMP (bottom panel). Each data point represents the mean ± SEM from at least three independent wells. (D) The requirement of extracellular calcium for MS4A1 ligand responses was assessed by stimulating HEK293 cells co-expressing GCaMP6s and either MS4A1 or mCherry with 2,5-DMP in the presence or absence of extracellular calcium. Data are presented as mean ± SEM from at least six wells per condition. ** p < 0.01, Tuckey’s test following one-way ANOVA compared to no extracellular calcium.

### Nitrogenous heterocyclic compounds activate MS4a1-expressing OSNs *in vivo*

These experiments were all carried out *in vitro*, and it remains unclear whether MS4A1 functions as an OR in an intact mouse. To test this, freely behaving mice were exposed to the *in vitro* identified ligands for MS4A1 in gas phase, and the activation of MS4A1-expressing cells was then assessed. 2,3-DMP and 2,5-DMP both activated MS4A1-expressing OSNs *in vivo* (Figures 6A and 6B). By contrast, ligands for other non-MS4A olfactory receptors, such as such as eugenol and CS_2_, did not activate MS4A1-expressing cells above background (Figures 6A and 6B). These experiments reveal that during conditions of active exploration, MS4A1-expressing cells respond to the chemicals we identified as MS4A1 ligands, indicating that MS4A1 functions as an olfactory receptor *in vivo*.

**Figure 6.**
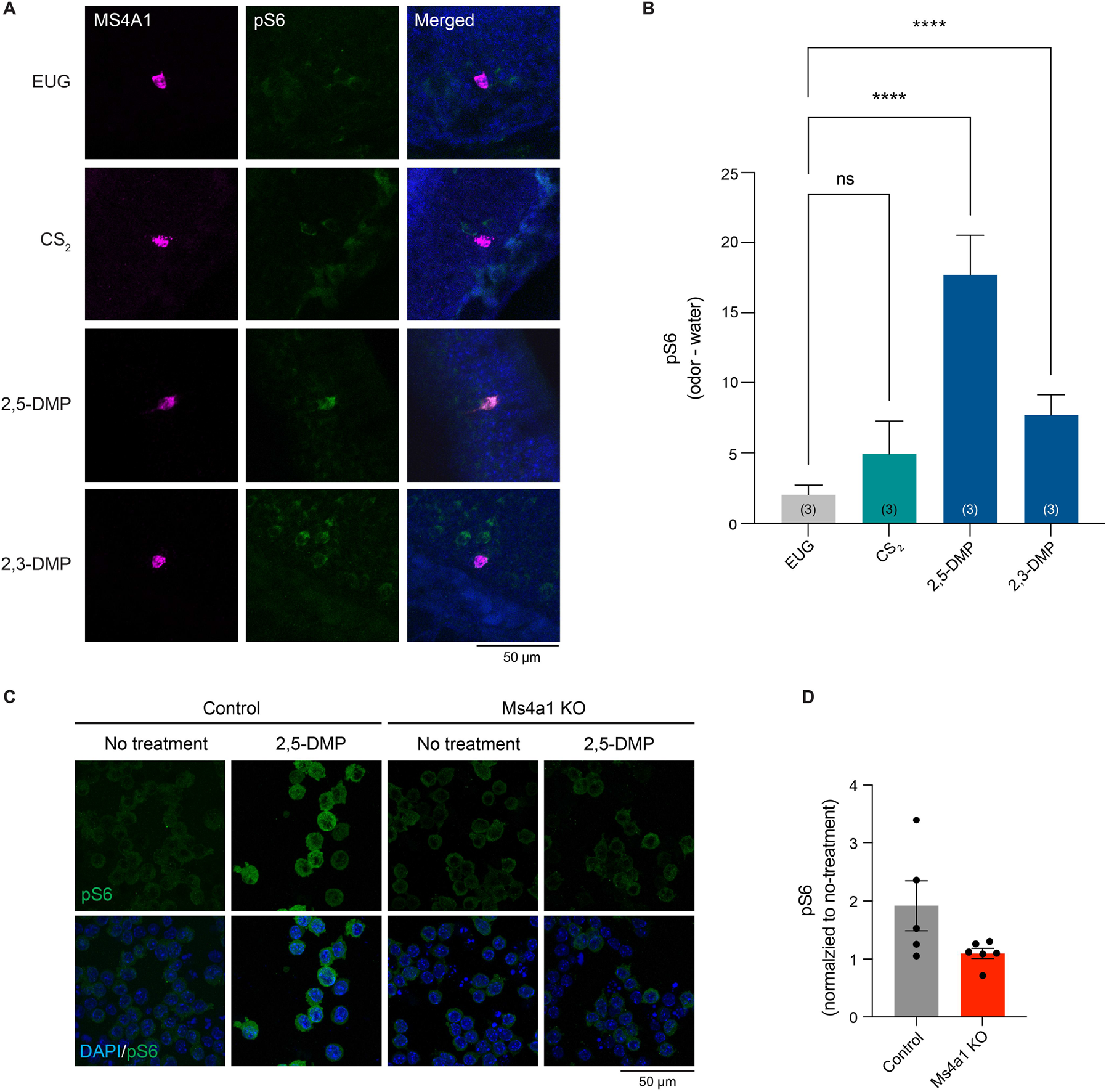
The MS4A1 ligands, 2,3-DMP and 2,5-DMP, activate MS4A1-expressing cells *in vivo*. (A) Example images of the main olfactory epithelia of mice exposed to the indicated odorant, immunostained for the neuronal activity marker, phospho-S6 (pSerine240/244) (green). *Ms4a1* is detected by fluorescent in situ hybridization (purple), see experimental procedures. EUG; eugenol, CS_2_; carbon disulfide, 2,3-DMP; 2,3-dimethylpyrazine, 2,5-DMP; 2,5-dimethylpyrazine. (B) Quantification of the proportion of pS6+ MS4A1-expressing OSNs (right panel) in response to the indicated odorants. Data are presented as mean ± SEM, from three independent experiments, **p < 0.01, ****p < 0.0001, Dunnett’s test following one-way ANOVA compared to null exposure. (C) Example images of 2,5-DMP stimulated wild-type (left panels) or *Ms4a1* knockout A20 cells, a BALB/c mouse B cell lymphoma cell line, immunostained for the activity marker phospho-S6 (pSerine240/244) (green). (D) Quantification of the proportion of pS6+ A20 cells in response to 2,5-DMP. Data are presented as mean ± SEM, from at least five independent experiments, (E)

## Discussion

Here, we took advantage of *Ms4a* knockout mice and used a combination of behavioral experiments and neuronal activation assays to show that MS4A proteins function as ORs within necklace subsystem OSNs *in vivo*. We also identified a new OSN subsystem that expresses MS4A1/CD20, but not other GPCR or MS4A ORs that were probed. MS4A1-expressing OSNs were sparse and located in the MOE, but were dispersed rather than geographically localized. We showed that MS4A1 recognizes nitrogenous heterocyclic compounds found in the urine of mouse predators and their sensing triggers innate, unlearned avoidance behavior.

These experiments convincingly demonstrate for the first time the existence of a non-GPCR family of mammalian ORs. All previously identified mammalian OR families were exclusively seven transmembrane spanning GPCRs (Buck and Axel, 1991). The discovery that *Ms4a* genes encode a polymorphic set of non-GPCR ORs raises questions about why this family of ORs evolved and what advantages it provides to mammalian olfaction. MS4A chemoreceptors respond to fairly non-descript chemical classes, including nitrogenous heterocyclic compounds, long-chain fatty acids, and steroids that can also be sensed by conventional GPCR ORs (Saito et al., 2009) (HCJ, SJP, and PLG unpublished observation). This finding suggests that MS4A proteins probably did not evolve to detect chemical compounds that the rest of the olfactory system doesn’t recognize, but rather, more likely evolved to mediate specific types of odor-driven behaviors. This is in line with the observation that in contrast to the “one-receptor, one-neuron” pattern of expression displayed by all other studied mammalian ORs, whereby each OSN expresses one, and only one of the approximately 1200 OR genes encoded by the murine genome (Buck and Axel, 1991; Mombaerts, 2004), many different MS4A proteins are co-expressed within the same necklace sensory neuron (Greer et al., 2016). This unorthodox pattern of expression suggests that MS4As may be important for mediating specific patterns of behavior, rather than for the exquisite discriminatory capacity, which the rest of the olfactory system possesses. Interestingly, our experiments suggest that necklace-expressed MS4A proteins, unlike MS4A1, which is expressed outside the necklace, do not play a major role in innate avoidance responses to their ligands. They likely are important in initiating other types of odor-driven behavior that remain to be defined. Perhaps the most likely behaviors induced by the necklace system may be social behaviors, since MS4A ligands are enriched for semiochemicals and pheromones (Greer et al., 2016). Moreover, necklace neurons have been implicated in the social transmission of food preference (Munger et al., 2010), a behavior whereby a mouse conveys its prior food experience to a conspecific animal. Thus, it is intriguing to speculate that MS4A proteins may participate in these or related behaviors.

While questions remain about the role of necklace-expressed MS4A members in odor-driven behaviors, this work identifies a function of the non-necklace cell-expressed MS4A family member, MS4A1. Here, we report that Ms4a1 encodes an olfactory chemoreceptor that is expressed in a previously uncharacterized population of OSNs within the MOE. MS4A1 detects nitrogenous heterocyclic compounds that are found at high abundance in the urine of natural predators of the mouse such as the wolf and the ferret, and we find that MS4A1 is required for the innate avoidance responses that mice exhibit in response to these compounds. Intriguingly, MS4A1 is expressed in a relatively small population of OSNs in the MOE, and prior work from a number of other laboratories has revealed that other discrete populations of sensory neurons, including TAAR-expressing cells, necklace cells, and Gruenberg ganglion neurons also mediate innate avoidance responses (Brechbühl et al., 2008; Dewan et al., 2013; Hu et al., 2007; Munger et al., 2010). Little is known about the neural circuitry downstream of these specialized OSN subpopulations that trigger innate avoidance responses, and in the future, elucidating how information flows from MS4A1-expressing neurons (and other olfactory subsystems that trigger innate avoidance), is likely to reveal how odor-driven innate avoidance behaviors are generated.

To address these questions, it will be important to fully characterize all of the different subpopulations of sensory neurons within the mammalian olfactory system. Our identification of a previously undescribed population of OSNs, with a corresponding newly characterized OR, suggests that there are likely still additional populations of OSNs (and ORs) to be found. The fact that MS4A1 falls outside of the canonical GPCR rubric, suggests that the traditional means of identifying additional ORs by relying on homology to known ORs, may be insufficient. RNA sequencing and spatial transcriptomics will facilitate the identification of additional olfactory subsystems.

This work may also have implications for understanding immune function. The only previously ascribed function for MS4A1 is as a co-receptor for the B-cell receptor in mature lymphocytes (Tedder and Engel, 1994; Tedder et al., 1988). The discovery that MS4A1 possesses chemoreceptive properties within the olfactory system suggests that MS4A1 may also function as a chemoreceptor in immune cells. Consistent with this idea, we find that B lymphocyte signaling is also activated by the MS4A1 ligand, 2,5-DMP (Figures 6C and 6D). Future work to identify what ligands MS4A1 senses in B lymphocytes and what effects their sensing has on B cell function are likely to be revealing. Other MS4A family members, besides MS4A1, are also found in other cell types and tissues throughout the body, including peripheral immune cells, microglia, reproductive cells, and lung cells (Hammond et al., 2019; Liu et al., 2019; Mattiola et al., 2019; Silva-Gomes et al., 2022). Polymorphisms in *Ms4a* genes have been strongly and reproducibly linked to a number of human diseases not obviously linked to olfaction, including Alzheimer’s Disease and asthma (Hollingworth et al., 2011; Lympany et al., 1992; Naj et al., 2011; Sandford et al., 1993). Therefore, characterizing the role of this family of chemoreceptors in non-olfactory contexts is likely to provide insight into organismal function in both healthy and disease states.

## Materials and Methods

### Mice

Mice were maintained under standard light/dark cycle conditions (12 hours light:12 hours dark) and were given food and water *ad libitum*. *Ms4a* cluster knockout mice were generated in the Datta Lab (Harvard Medical School) by standard approaches using CRISPR/Cas9 technology and will be described in detail elsewhere. *Ms4a6c* knockout mice were generated by KOMP using homologous recombination. These *Ms4a* knockout mice were kindly provided by the Datta Laboratory. *Ms4a1* knockout mice (C57/BL6) were obtained from the Tedder Lab, Duke University (Uchida et al., 2004). All behavioral and immunostaining experiments with knockout mice were performed with littermate wild-type control mice. The following primer sequences were used for genotyping: 1) *Ms4a* cluster knockout, common primer (5’-GACAAATGAACTAACCTTGCTTGG-3’), wild type specific primer (5’-TCCAGTGGAAGTGGTTTTGC-3’), and deletion specific primer (5’-GCCTTGGCTAGGCTACAACC) were used to amplify a fragment of 412 bp from the wild-type allele and 259 bp from the deleted allele. 2) *Ms4a6c* knockout, a 204 bp fragment from the wild type allele was amplified with one primer set (5’-GGACAGAAACGCCTAAAGGT-3’ and 5’-AGAGAAGGGAGATGGTGACTACTA-3’), and a second set of primers (5’-CTAAACTCAAGAGGTCATTGAAG-3’ and 5’-GCAGCGCATCGCCTTCTATC-3’) amplified a 280 bp fragment from the targeted allele. 3) *Ms4a1* knockout, a 487 bp fragment from the wild type allele was amplified with one primer set (5’-GATATCTACGACTGTGAACC-3’ and 5’-TGGCATGTGCCAGTAAGCC-3’), and a second set of primers (5’-TTTGGGGGCTGTCCAAATCATG-3’ and 5’-CATCGCCGACAGAATGCCC-3’) amplified a 445 bp fragment from the targeted allele. All mouse husbandry and experiments were performed following institutional and federal guidelines and approved by University of Massachusetts Chan Medical School’s Institutional Animal Care and Use Committee.

### Plasmids

mCherry was cloned into the pcDNA3.1(+) backbone (mCherry-pcDNA3.1). The complete coding sequences of mouse Ms4a2, Ms4a3, Ms4a4a, Ms4a4b, Ms4a4c, Ms4a4d, Ms4a5, Ms4a6b, Ms4a6c, Ms4a7, Ms4a8a, Ms4a10, Ms4a13, Ms4a15, Ms4a18 were cloned into mCherry-pcDNA3.1. Mouse or human Ms4a1 DNA coding sequences were cloned into the tetracycline inducible mammalian expression plasmid, pcDNA5-FRT-TO. pGP-CMV-GCaMP6s was a gift from Douglas Kim (Addgene, #40753). pISH-Gucy1b2-probe1 (Addgene, #105454), pISH-Gucy1b2-probe2 (Addgene, #105455), pISH-Trpc2-probe1 (Addgene, #105473), pISH-Trpc2-probe2 (Addgene, #105474), pISH-Trpm5 (Addgene, #105993), pISH-Gucy2d-1 (Addgene, #105459), pISH-V1rb1 (Addgene, #16010) were gifts from Peter Mombaerts.

### Antibodies

Primary antibodies/concentrations used were as follows: rabbit anti-phospho-S6 ribosomal protein (Serine240/244) (1:100, Cell Signaling Technologies, #2215), rabbit anti-phospho-S6 ribosomal protein (Serine244/247) (1:150, Invitrogen, #44-923G), rabbit anti-MS4A1/CD20 (1:250 for immunostaining, 1:100 for iDISCO, Cell Signaling Technology, #98708), rabbit anti-MS4A1 (1:200, MyBioSource, #MBS2051903), goat anti-MS4A1 (1:50, Santa Cruz Biotechnology, #sc-7735), rat anti-MS4A1 (1:100, LifeSpan Biosciences, #LS-C107163-100), guinea pig anti-VGLUT2 (1:500, SYSY, #135 404), rabbit anti-NeuN (1:500, Abcam, #ab104225), rabbit anti-KI18 (1:500, Abcam, #ab52948), rabbit anti-KI17 (1:500, Abcam, #ab53707), goat anti-NeuroD1 (1:50, R&D Systems, #AF2746), goat anti-OMP (1:1000, Wako Chemicals, #544-10001-WAKO), rabbit anti-CNGA2 (1:200, Alomone Labs, #APC-045), and rabbit anti-PDE2A (1:500, FabGennix, #PD2A-101AP).

Secondary antibodies/concentrations used were as follows: Alpaca anti-rabbit-Alexa488 (1:333, Jackson Immunoresearch, #611-545-215), alpaca anti-rabbit-rhodamine red X (RRX) (1:333, Jackson Immunoresearch, #611-295-215), goat anti-rabbit-Alexa647 (1:333, Invitrogen, #A-21245), bovine anti-goat Alexa488 (1:333, Jackson Immunoresearch, #805-545-180), bovine anti-goat Alexa647 (1:333, Jackson Immunoresearch, #805-605-180), goat anti-rat Alexa488 (1:333, Invitrogen, #A-11006), donkey anti-rat RRX (1:333, Jackson Immunoresearch, #712-295-153), and goat anti-guinea pig Alexa647 (1:333, Invitrogen, #A21450).

### Odors

Eugenol, CS_2_, 2,3-DMP, 2,5-DMP, OA, ALA, indole, quinoline, pyridine, pyrrolidine, vanillin, and isoamyl alcohol (IAA) were purchased from Sigma-Aldrich. TMT was purchased from BioSRQ. All odor compounds were obtained at the highest purity possible.

### Odor exposure for phospho-S6 immunostaining

8–12-week-old mice, including *Ms4a* cluster knockout, *Ms4a6c* knockout, *Ms4a1* knockout, and littermate wild-type control mice, were individually acclimated to the clean plastic cage (Innovive, # M-BTM) for at least 16 hours before the start of experiments. Before introducing odors to the mice, the mice were fasted for 2 hours. To initiate the experiments, water or odor stimulus (eugenol, CS_2_, 2,3-DMP, 2,5-DMP, OA, ALA) were introduced into each cage. The stimuli were applied by placing 150 µL of water or odorant on filter paper (Sigma-Aldrich, #WHA10347509) in 35 mm petri dishes. After 2 hours of exposure to the odor, the mice were euthanized, and nasal epithelial sections were collected.

### Tissue slice preparation

The mice were euthanized, and their noses, including the olfactory epithelia and attached olfactory bulbs, were dissected from the skull. The dissected tissue was fixed overnight in 4% paraformaldehyde (PFA, Electron Microscopy Sciences, #15714) in phosphate-buffered saline (PBS) at 4°C. After washing three times for 5 minutes with 1X PBS, noses were decalcified overnight at 4°C in 0.45M EDTA in PBS. Subsequently, the noses were sequentially incubated in 10%, 20%, and 30% sucrose in PBS (Sigma-Aldrich, #S0389) overnight at 4°C. Finally, the tissues were embedded in Tissue Freezing Medium (Tissue-Tek, #4583). Cryosections of 20 micron thickness were cut onto Superfrost Plus glass slides (VWR #48311-703) and stored at -80°C until staining.

### Combined pS6 immunostaining and RNAscope fluorescent in situ hybridization (FISH)

For *Car2* and *Ms4a1* RNA detection, RNAscope FISH was performed on nasal epithelial sections from 8-12-week-old C57/BL6 wild-type, *Ms4a* cluster knockout, *Ms4a1* knockout, and *Ms4a6c* knockout mice that were exposed to water or odors as described above. The staining protocol followed the guidelines provided in the Advanced Cell Diagnostics RNAscope Multiplex Fluorescent Reagent Kit v2 Assay User Manual (323100-USM) without the target retrieval step, and all required reagents were obtained from RNAscope Multiplex Fluorescent Detection kit v2 (Advanced Cell Diagnostics, #323110).

Frozen sections were thawed at room temperature for 10 minutes, then were fixed with 4% PFA in PBS for 15 minutes at 4°C. The sections were dehydrated with 50%, 70%, and 100% ethanol for 5 minutes each at room temperature (RT). The sections were treated with hydrogen peroxide for 10 minutes at RT, then washed with MilliQ water three times. Sections were treated with protease III at 40°C in a hybridization oven (HybEZ oven, Advanced Cell Diagnostics, #310010) for 30 minutes, then washed with MilliQ water three times. Subsequently, the sections were hybridized with either the 3-Plex positive control RNA probe, the 3-Plex negative control RNA probe, Car2 RNA probe, or Ms4a1 RNA probe in a 1:50 ratio for 2 hours at 40°C in the oven.

To amplify hybridization signal, the section slides underwent incubation with three different amplifiers: AMP1 for 30 minutes, AMP2 for 30 minutes, and AMP3 for 15 minutes, all at 40°C in the oven. After the amplification steps, slides were treated with horseradish peroxidase (HRP) for 15 minutes at 40°C in the oven. Following this, the sections were incubated with diluted TSA plus Cy-3 (1:750, PerkinElmer, #NEL741001KT) for 30 minutes at 40°C in the oven, then the sections were incubated with HRP blocker for 15 minutes at 40°C in the oven. Washing was performed with RNAscope wash buffer (2 minutes twice at RT) between each step following probe hybridization. The mouse target probes used in this study were as follows: Ms4a1-c2, #318671-C2 and Car2-c2, #313781-C2.

For the pos-RNAscope immunostaining, sections were blocked with blocking buffer (0.1% Triton X-100 (Sigma-Aldrich, #X100) 5% Normal Donkey serum (Jackson ImmunoResearch, #017-000-121), 3% Bovine Serum Albumin (VWR, #97061-416) in PBS) for 30 minutes at RT. Sections were then incubated with anti-pS6 antibodies (1:100) in blocking buffer overnight at 4°C. On the following day, the slides were washed three times with PBS (5 minutes each at RT) and then incubated with the secondary antibody (1:300) in blocking solution for 45 minutes at RT. Afterwards, the slides were washed three times with PBS (5 minutes each at RT) and mounted using Vectashield antifade mounting media with DAPI (Vector Laboratories, #H-1200-10). To secure the coverslips, nail polish was applied, and the slides were imaged using confocal microscopy, following the procedures described below.

### Conventional FISH

For Gucy1b2, Trpc2, Trpm5, Gucy2d, and V1rb1 RNA detection, conventional FISH was performed on nasal epithelial sections from C57/BL6 wild-type mice, following a modified protocol (Ishii et al., 2004). The RNA probes for these genes were previously described (Omura and Mombaerts, 2014, 2015; Parrilla et al., 2016; Rodriguez et al., 2002). Fluorescein isothiocyanate (FITC, Roche, #11685619910)-labeled riboprobes were generated through *in vitro* transcription from fully linearized and purified template plasmids (described in Plasmids) containing target gene sequences, using equilibrated phenol (Sigma-Aldrich, #P9346) – chloroform/isoamyl alcohol (Sigma-Aldrich, #25666). Frozen sections were air-dried, fixed with 4% PFA/1X PBS for 10 minutes, and then acetylated with a mixture of 0.1M triethanolamine (Sigma-Aldrich, #90279) and 0.25% acetic anhydride (Sigma-Aldrich, #320102) for 15 minutes at RT. Pre-hybridization was performed in a prehybridization solution (10 mM Tris, pH 7.5, 600 mM NaCl, 1 mM EDTA, 0.25% SDS, 1X Denhardt’s (Sigma, #D-2532), 50% formamide (Roche, #1814320), 300µg/ml yeast tRNA (Sigma-Aldrich, #R6750)) for 5 hours at 65 °C. Following pre-hybridization, the sections were hybridized overnight at 60 °C with FITC-labeled RNA probes (1 µg/ml) in a hybridization solution (10 mM Tris, pH 7.5, 200 mM NaCl, 5 mM EDTA, 0.25% SDS, 1X Denhardt’s, 50% formamide, 300 µg/ml yeast tRNA, 10% dextran sulfate (Bio Basic, #DB0160), 5 mM NaH_2_PO_4_, 5 mM Na_2_HPO_4_).

On the subsequent day, the slides were sequentially washed with the following buffers: 5X standard saline citrate buffer (Invitrogen, #AM9765) (10 minutes, 65 °C), 50% formamide/1X SSC (30 minutes, 65 °C), TNE buffer (10 mM Tris, pH 7.5, 0.5M NaCl, 1 mM EDTA) (20 minutes, 37 °C), 2X SSC (20 minutes, 65 °C), and 0.2X SSC (20 minutes, 65 °C) twice. Quenching of endogenous peroxidase activity was performed using 1% H_2_O_2_ (Sigma-Aldrich, #216763)/1X PBS for 15 minutes at 4 °C, followed by blocking with a blocking buffer (0.1 M Tris, pH 7.5, 100 mM maleic acid (Sigma-Aldrich, #M0375), 150 mM NaCl, 0.1% Tween-20, 2% blocking reagent (Roche, #11096176001), 10% heat-inactivated sheep serum (Equitech-Bio, #SS32-0100)) for 30 minutes at RT. The samples were then incubated overnight at 4°C with anti-fluorescein-POD (1:2000, Roche, #11426346910).

On the third day, the slides were washed three times with PBST (1X PBS/0.1% Tween-20), incubated with diluted TSA plus fluorescein (1:50) for 5-10 minutes at RT, and washed five times with PBST. Finally, the sections were immunostained with anti-MS4A1 antibodies and imaged using confocal microscopy, following the procedures described below.

### Immunostaining

Sections were incubated in a blocking solution containing 5% normal donkey serum, 0.1% Triton-X100, and 1X Tris-buffered saline (TBS) for 1 hour at RT. Subsequently, sections were incubated overnight at 4°C with primary antibodies diluted in blocking solution. On the following day, slides were washed three times with TBST (0.1% Triton-X100 in TBS) and then incubated with secondary antibodies in blocking solution for 1 hour at RT. Afterwards, the slides were washed three times with TBST, counterstained, and mounted using Vectashield antifade mounting media with DAPI. To secure the coverslips, nail polish was applied, and the slides were imaged using confocal microscopy, following the procedures described below.

### Confocal microscopy

Slides were imaged using an LSM 900 Airyscan2 confocal microscope (Zeiss) equipped with various objective lenses, including 10X/0.45 M27, 20X/0.8 M27, 40X/1.1 water Corr M72, and 63X/1.4 oil DIC. To enhance image quality, acquired digital images were processed by applying a median filter to remove debris that was significantly smaller than the structures being analyzed. Additionally, multi-channel Z-stacks were projected into two dimensions using Zen blue 3.1 software (Zeiss).

### Quantification of phospho-S6 (pS6) positive cells

Car2 or Ms4a1 positive cells were selected using imageJ sorftware. pS6 intensity was determined following subtraction of background signal from cells in the olfactory epithelium lacking Car2 or Ms4a1 signal. For necklace cells, 12 sections from the posterior olfactory epithelium were collected, and three Car2-positive regions were randomly selected from each section to perform quantification. For Ms4a1-positive cells, 24 sections equally spaced throughout the anterior to posterior axis of the olfactory epithelium were collected, and all the Ms4a1-positive cells from these sections were analyzed. Analysis was performed blinded to genotype and stimulus to ensure unbiased quantification.

### iDISCO

iDISCO was performed on the olfactory bulbs of 8-12 week old C57/BL6 wild-type mice following the protocol described by Renier and colleagues (Renier et al., 2014) (online protocol: http://idisco.info/idisco-protocol). Initially, mice were euthanized, and their olfactory bulbs were dissected and fixed for 2 hours in 4% PFA in PBS at 4 °C. The samples were then washed three times with PBS for 30 minutes each at RT and then dehydrated using a series of methanol solutions (20%, 40%, 60%, 80%, 100%, Sigma-Aldrich, #179337-4X4L) for 1 hour each at RT. Subsequently, the samples were incubated overnight in a mixture of 66% dichloromethane (Sigma-Aldrich, #270997) and 33% methanol at RT and washed twice with methanol. The samples were then bleached using 5% H_2_O_2_/ methanol solution at 4 °C overnight. Following bleaching, the samples were rehydrated using a series of methanol solutions (80%, 60%, 40%, 20%) and PBS for 1 hour each at RT.

The samples were incubated in permeabilization solution (0.2% Triton X-100, 0.3 M glycine (Merck, #G5417), 20% DMSO (Corning, #25-950-CQC) in PBS) at 37 °C overnight and then placed in blocking solution (0.2% Triton X-100, 6% donkey serum, 10% DMSO in PBS) at 37 °C overnight. Subsequently, the samples were washed in PTwH buffer (0.2% Tween-20, 10 μg/ml heparin (Sigma-Aldrich, #H3393) in PBS) overnight and incubated with primary antibodies diluted in PTwH buffer supplemented with 5% DMSO and 3% donkey serum at 37 °C for 3 days. The samples were extensively washed in PTwH buffer for 1 day and then incubated with secondary antibodies diluted in PTwH buffer with 3% donkey serum at 37 °C for 3 days. Finally, the samples were washed in PTwH buffer overnight before the clearing and imaging process. For clearing, the bulb samples were dehydrated using a series of methanol solutions (20%, 40%, 60%, 80%, 100%, 100%) for 1 hour each at RT. The samples were incubated in 66% dichloromethane/33% methanol at RT for 3 hours and then incubated in 100% dichloromethane until they sank to the bottom of the vial. Finally, the samples were incubated in dibenzyl ether (Sigma, #108014) at RT overnight. The cleared samples were directly imaged using a light-sheet microscope (Ultramicroscope II; LaVision BioTec). The images were acquired using InspectorPro software (LaVision BioTec), and three-dimensional reconstruction and analysis were performed using Imaris x64 software (v.8.0.1, Bitplane).

### Odor-driven behavior assay

For behavioral experiments, 8–12-week-old *Ms4a* cluster knockout, *Ms4a6c* knockout, *Ms4a1* knockout and littermate wild-type control mice were group housed in the behavioral assay room and allowed to acclimate for at least one day prior to the start of the experiments. Two hours prior to the behavioral assay, mice were individually fasted in their home cage. During the experiment, mice were placed in single use, disposable cages (Innovive, #M-BTM) with a disposable paper curtain separating the avoidance zone from the odorized zone. Only the odorized zone was enclosed by an acrylic sheet on the top of cage. A clean filter paper was placed in a 35 mm petri dish within the odorized zone. Without any odor stimulus, mice were first allowed to freely explore their surroundings for 30 minutes. Subsequently, the mice were exposed to water (40 µL) applied to the filter paper in the odorized zone for a duration of 3 minutes. After water exposure, the same mouse was then exposed to 40 µL of odorant for 3 minutes. The odorant was delivered onto a fresh filter paper for the experiment. Animal behavior during the entire experiment, including habituation, water exposure, and odor exposure was recorded with a webcam (Logitech, #LOWCC920S). Those videos were then analyzed with ezTrack (Pennington et al., 2019).

### Elevated plus maze assay

The elevated plus maze (EPM) apparatus consisted of plus-shaped (+) apparatus with two open and two enclosed arms. The closed arms are enclosed by black walls (30 × 5 × 15 cm) and the open arms are exposed (30 × 5 × 0.25 cm). The maze was elevated 45 cm above the floor, and a red fluorescent light was positioned 1 meter above the maze as light source. The whole assay was performed in a darkroom. 8–12-week-old *Ms4a* cluster knockout, *Ms4a6c* knockout, *Ms4a1* knockout, and littermate wild-type control mice were group housed in the darkroom for 1 hour prior to the experiment. Individual mice were placed at the center of the maze, and the mouse was allowed to freely explore the maze for 5 minutes. The time the mice stayed in the open arm and the closed arm is automatically measured by the system. The maze was cleaned with rubbing alcohol prior and between experiments. The maze was design by Andrew Tapper (Molas et al., 2017).

### RNA-Sequencing

Three 8-12-week old *Ms4a* cluster knockout, *Ms4a6c* knockout, and littermate wild-type control mice were euthanized according to our IACUC protocol, and the main olfactory epithelium was dissected and immersed quickly in 0.6 ml of ice-cold Trizol (Invitrogen, #15596018). Olfactory epithelia were ground and homogenized with nuclease-free disposable pestles (Fisher Scientific, #12-141-368) in an Eppendorf tube (Eppendorf, #2231000347). The homogenized sample was incubated for 5 minutes at room temperature. 0.1 µL of chloroform was added to the sample and mixed by inverting for 20 seconds. The sample was then incubated for 3 minutes at RT. The sample was centrifuged at 12,000 xg for 15 minutes at 4 °C. The clear aqueous phase was collected into a nuclease-free Eppendorf tube, and an equal volume of 70% ethanol (Sigma-Aldrich, #E7023) was added and mixed. The sample was then loaded into RNeasy spin column and RNA was extracted with a RNeasy mini kit (Qiagen, #74104). NEBNext Ultra II Directional RNA Library Prep Kit for Illumina (NEB, #E7490S and #E7760S) was used to construct sequencing libraries following the manufacturer’s guidelines with one alteration, which was to increase the insert length to around 300 bp. Libraries were sequenced using paired-end 150 cycle reads on a Novaseq6000 (Novogene). The sequencing reads were processed using the DolphinNext RNAseq pipeline (https://dolphinnext.umassmed.edu/index.php?np=1&id=732) (Yukselen et al., 2020). The default settings were employed, except that STAR v.2.6.1 and RSEM v.1.3.1 were used for alignment and quantification, respectively (Dobin et al., 2013; Li and Dewey, 2011). Transcriptome build: gencode M25. The count matrix was loaded to R (v.4.0.0 or later), and DEseq2 (v.1.30.1) was used to normalize the matrix and perform differential gene expression analysis (Love et al., 2014).

### Cell culture and transfection

Human embryonic kidney 293 (HEK293) cells (ATCC, #CRL-3216) were cultured in complete media, which consisted of DMEM-high glucose (Sigma-Aldrich, #D5671) supplemented with 10% heat-inactivated fetal bovine serum (R&D Systems, #S12450H), penicillin/streptomycin/glutamine (Gibco, #10378016), and maintained in a 5% CO_2_ humidified tissue culture incubator at 37 °C. For calcium imaging experiments, HEK293 cells were transfected with calcium phosphate. 0.5 x 10^6^ cells were seeded in each well of a 12-well plate, and 1 µg of GCaMP6s and 1 µg of either mCherry or mCherry-MS4A fusion plasmid DNA were mixed with 250 mM CaCl_2_. The solutions were resuspended by pipetting four times and combined with 2X HBS (containing 50 mM HEPES, 10 mM KCl, 12 mM D-glucose, 280 mM NaCl, and 1.5 mM Na_2_PO_4_ at pH 7.06). The reaction mixtures were incubated for 5 minutes at RT and then added dropwise to each well. For the expression of MS4A1 protein, 1 hour after transfection, tetracycline (Sigma-Aldrich, #T7660) was added to a final concentration of 1 µg/mL. For surface immunostaining, HEK293 cells were plated on 12 mm round German glass coverslips (Bellco Biotechnology, #1943-10012A) coated with poly-d-lysine hydrobromide (Sigma-Aldrich, #P0296) in a 24-well plate and incubated in complete media. The cells were transfected using the same calcium phosphate method as described above.

### Generation of lentiviral CRISPR/CAS9-mediated Ms4a1 knockout B cell lines and pS6 immunostaining

To produce lentivirus, HEK293T cells were transfected with pLentiCRISPR v2 plasmids containing guide sequences targeting the control gene (ACTATCATGGCACCCAATTG) and the Ms4a1 gene (GATGGGTGCGAAGACCCCTG) (Wang et al., 2014), along with delta-Vpr packaging plasmid and the VSV-G envelope plasmid. X-tremeGENE 9 Transfection reagent (Roche, #XTG9-RO) was used for the transfection. Lentivirus-containing media were collected 24 hours after changing to fresh media. The supernatant containing the virus was used without concentration after one cycle of freeze and thaw to eliminate any residual HEK293T cells. The virus was then transduced into A20 cells (ATCC, #TIB-208), a mouse BALB/c B cell lymphoma line (Kim et al., 1979), with 10 µg/ml polybrene for 1 hour using centrifugation at RT. The cells were incubated for 24 hours, and the media was replaced with media containing 1 µg/ml puromycin to select for transduced cells. Single-cell clones were selected, expanded, and subjected to Western blot analysis to generate single cell-derived Ms4a1 knockout A20 cells. For the pS6 activation experiments, both control and Ms4a1 knockout cells on coverglass were incubated in serum-free media for 30 minutes and treated with either vehicle or 2,5-DMP for 30 minutes in a tissue culture incubator at 37 °C. Subsequently, the cells were fixed with 4% PFA/PBS for 20 minutes at RT, followed by three washes with PBS. The cells were incubated in blocking buffer for 30 minutes at RT, and then with anti-pS6 antibodies (1:400) in blocking buffer overnight at 4°C. On the following day, the cells were washed three times with PBS and then incubated with secondary antibody (1:300) in blocking solution for 45 minutes at RT. After three washes with PBS, the cells were mounted using Vectashield antifade mounting media with DAPI.

### Calcium imaging

24 hours post-transfection, media in the wells were aspirated and washed twice with 1X Ringer’s solution supplemented with 1 mM CaCl_2_ (Ca^2+^-Ringer’s solution). The wells were then incubated for 30 minutes in the cell culture incubator with Ca^2+^-Ringer’s solution. Following the incubation, the plate was transferred to a Lionheart LX Automated microscope (BioTek), and calcium imaging was performed using Gen5 software (BioTek). Preliminary images were acquired with brightfield, RFP, and GFP filters at 20X magnification prior to each experiment, focusing on an imaging field containing cell numbers between 50 and 200. Subsequently, using the same field of view and fixed z-axis, images were captured for 10 minutes (1 FPS). During the kinetic image acquisition, either Ca^2+^-Ringer’s solution as a negative control or specific odorants (50 µM for 2,3-DMP, 2,5-DMP, 2,6-DMP, indole, quinoline, pyridine, pyrrolidine, vanillin, IAA) solubilized in Ca^2+^-Ringer’s solution were pipetted into the upper edge of each well after 360 seconds for a duration of 10 seconds. To determine dose-response curves and calculate the EC50, HEK293 cells co-expressing GCaMP6s and either mCherry or mCherry-MS4A1 were treated with six logarithmic orders of 2,3-DMP or 2,5-DMP (ranging from 10 nM to 1 mM) starting with the lowest concentration. For experiments conducted without extracellular calcium, all solutions were replaced with 1X Ringer’s solution supplemented with 1 mM EGTA and 1 mM EDTA to chelate calcium. All acquired images were aligned to the first image of each experiment using Gen5 software, and subsequent images were analyzed using Fiji software.

### Analysis of calcium imaging data

Mp4 videos were converted into a sequence of PNG images with ffmpeg software, the image sequence was then imported into Fiji. GCaMP6s and mCherry positive cells were selected, and their GCaMP6s intensities were calculated across the whole image sequence, which was subsequently analyzed using a customized R script. Briefly, for each selected cell, the average intensity and standard error of GCaMP6s 30 seconds prior to ligand presentation was calculated. 2.5-fold of the standard error above mean intensity was then used as a threshold to determine if the cell responded to the odor.

### Statistics and reproducibility

For quantification of pS6, at least three biological repeats were performed for each odorant treatment. All analyses were conducted blinded to genotype and stimulus to ensure unbiased quantification. One-way ANOVA was performed to calculate the statistical difference of mean intensity of pS6 from all groups. A post-hoc Dunnett’s test was used to determine if the mean intensity of pS6 from a given odorant treatment was significantly different from the eugenol control group.

For the odor-driven behavior assay, at least five biological repeats were performed for each odorant. The analysis was conducted in an automated manner whenever possible in the absence of human supervision to ensure blinding to genotype and stimulus. For the total distance traveled, an unpaired Welch t-test was performed to calculate statistical differences. For the distance between the mouse’s center of mass and the odor, a paired t-test was performed to calculate statistical differences. For comparing the proportion of time spent in the odorized zone and the distance between the mouse’s center of mass and odor (unpaired), a one-way ANOVA was performed to calculate statistical differences between all groups. A post-hoc Dunnett’s Test was applied to determine if the values from a given odorant treatment are significantly different from those from the water control group. For the avoidance index, an unpaired Welch t-test was used to compare each knockout group and their wild-type littermates. A false-discovery rate was controlled using a two-stage step-up developed by Benjamini, Krieger, and Yekutieli.

For the EPM assay, at least five biological repeats were performed for each genotype. An unpaired Welch t-test was used to compare time spent in the open arm between groups.

For calcium imaging, at least nine biological repeats were performed for each odorant. The analysis was conducted blinded to protein expressed and stimulus to ensure unbiased quantification. For identifying 2,5-DMP responsive MS4A receptors, a one-way ANOVA was performed using a post-hoc Dunnett’s test. For screening chemicals that might activate MS4A1-expressing cells, a one-way ANOVA was performed with a post-hoc Dunnett’s test to compare to cells exposed to Ringer’s solution only. For assessing whether the presence of extracellular calcium affects the response rate of MS4A1-expressing cells, a one-way ANOVA was performed with a post-hoc Tukey’s test.

For all the symbols indicating statistical significance in this article: ****, p<0.0001; ***, p<0.001; **, p<0.01; *, p<0.05; ns, p ≥ 0.05.

## Data availability

All RNA sequencing data described in this manuscript are deposited at GEO accession GSE240378, which is associated with Figures S1B and S2B.

## Code availability

All the scripts used for this study can be found at: https://github.com/Greerlab/CD20_2023_paper.

## Acknowledgements

We thank Judy Lieberman, Eric Greer, Jin Zhang, Bob Datta, and members of the Greer Lab for helpful comments on the manuscript. We would like to thank Bob Datta and Thomas Tedder for generously providing mouse lines, Rubing Zhao-Shea for technical assistance with the elevated plus maze experiments, and Namgyu Lee and Leonid Yurkovetskiy for helping to generate Ms4a1 knockout A20 cell lines. P.L.G. was supported by fellowships from the Smith Family Foundation, the Searle Scholars Program, the Rita Allen Foundation, the Whitehall Foundation, and by grants DP2 OD027719-01 and NIH 5 KL2 TR001455-04 from the National Institutes of Health. H.C.J and I.H.W were supported by Mello Fellowships. S.J.P was supported by Mogam Science Fellowship. D.M.B is a Biogen Fellow of the Life Sciences Research Foundation and an Interdisciplinary Scholar of the Wu Tsai Neurosciences Institute at Stanford University.

