## Supplementary Information for "CD20 is a mammalian odorant receptor expressed in a subset of olfactory sensory neurons that mediates innate avoidance of predators"

**Figure S1 related to Figure 1.**
(A) PCR analysis of genomic DNA from wild-type and heterozygous or homozygous Ms4a cluster knockout mice.
(B) Heatmap of expression of Ms4a family member mRNA transcripts detected by RNA sequencing experiments performed on RNA isolated from the main olfactory epithelia of wild-type and Ms4a cluster knockout mice.
(C) Quantification of the velocity of Ms4a cluster knockout mice and their wild-type littermate controls during the odor avoidance assays.
(D) Heat maps (top panels) of the occupancy of wild-type mice in the odor avoidance chamber in response to the indicated odorants. Small square represents location of odorant, and dashed line demarcates the odor avoidance zone from the rest of the chamber. Scale bar, 5 cm. Bottom panels represent quantification of the avoidance mice exhibit in response to the indicated odors. **$p < 0.01$, ***$p < 0.001$, a paired t test compared to water. EUG; eugenol, LA; linolenic acid, OA; oleic acid, ALA; alpha-linolenic acid, AA; arachidonic acid, 2,3-DMP; 2,3-dimethylpyrazine, 2,5-DMP; 2,5-dimethylpyrazine.
(E) Quantification of the amount of time male (left panel) and female (right panel) Ms4a cluster knockout mice and their wild-type littermate controls spend in open arms in an elevated plus maze assay. Unpaired Welch t-test compared to wild-type mice.

**Figure S2 related to Figure 2.**
(A) PCR genotyping analysis of the wild-type (top) and deleted (bottom) *Ms4a6c* alleles from, wild-type and heterozygous or homozygous *Ms4a6c* knockout mice.
(B) Heatmap of expression of Ms4a family member mRNA transcripts detected by RNA sequencing experiments performed on RNA isolated from the main olfactory epithelia of wild-type and *Ms4a6c* knockout mice. The residual *Ms4a6c* transcripts detected in *Ms4a6c* knockout mice all map to 5' UTR regions of the gene, which were not targeted.
(C) Quantification of the velocity of *Ms4a6c* knockout mice and their wild-type littermate controls during the odor avoidance assays. Unpaired Welch t-test compared to wild-type mice.
(D) Quantification of the amount of time male (left panel) and female (right panel) *Ms4a6c* knockout mice and their wild-type littermate controls spend in open arms in an elevated plus maze assay. Unpaired Welch t-test compared to wild-type mice.

**Figure S3 related to Figure 3.**
(A) Representative confocal images of HEK293 cells transfected with an N-terminal mCherry-fusion version of MS4A1 (purple) in which the surface expression of MS4A1 was determined using non-permeabilized staining conditions and an anti-MS4A1 antibody (cyan).
(B) Quantification of the velocity of *Ms4a1* knockout mice and their wild-type littermate controls during the odor avoidance assays. Unpaired Welch t-test compared to wild-type mice.
(C) Quantification of the amount of time male (left panel) and female (right panel) *Ms4a1* knockout mice and their wild-type littermate controls spend in open arms in an elevated plus maze assay. Unpaired Welch t-test compared to wild-type mice.

(D) Quantification of the velocity of *Rag1* knockout mice and their wild-type littermate controls during the odor avoidance assays. Unpaired Welch t-test compared to wild-type mice.
(E) Quantification of odor avoidance behavior. The distance between the average center of mass of the mouse and the location of odorant was determined for *Rag1* knockout mice (right). Each circle represents an individual mouse. Data are presented as mean ± SEM, n ≥ five independent experiments, $^{*}p < 0.05$, a paired t test compared to water exposure. 2,5-DMP; 2,5-dimethylpyrazine.

**Figure S5 related to Figure 5.**
(A) Quantification of baseline responses of HEK293 cells expressing MS4A1 protein in the absence of chemical stimulation. n ≥ six independent experiments.
(B) Quantification of responses of HEK293 cells expressing either mouse MS4A1 protein or human MS4A1 protein in response to 2,5-DMP. n ≥ 6 independent experiments. $^{**}p < 0.01$, Dunnett's test following one-way ANOVA compared to mCherry alone. 2,5-DMP; 2,5-dimethylpyrazine.
(C) Quantification of responses of HEK293 cells expressing mCherry or MS4A1 to the indicated chemicals as in Figures 5A and 5B. n ≥ three independent experiments. $^{*}p < 0.05$, $^{**}p < 0.01$, $^{***}p < 0.001$, a paired t test compared to GCaMP6s only. 2,3-DMP; 2,3-dimethylpyrazine, 2,6-DMP; 2,6-dimethylpyrazine.

**Figure S1 related to Figure 1**

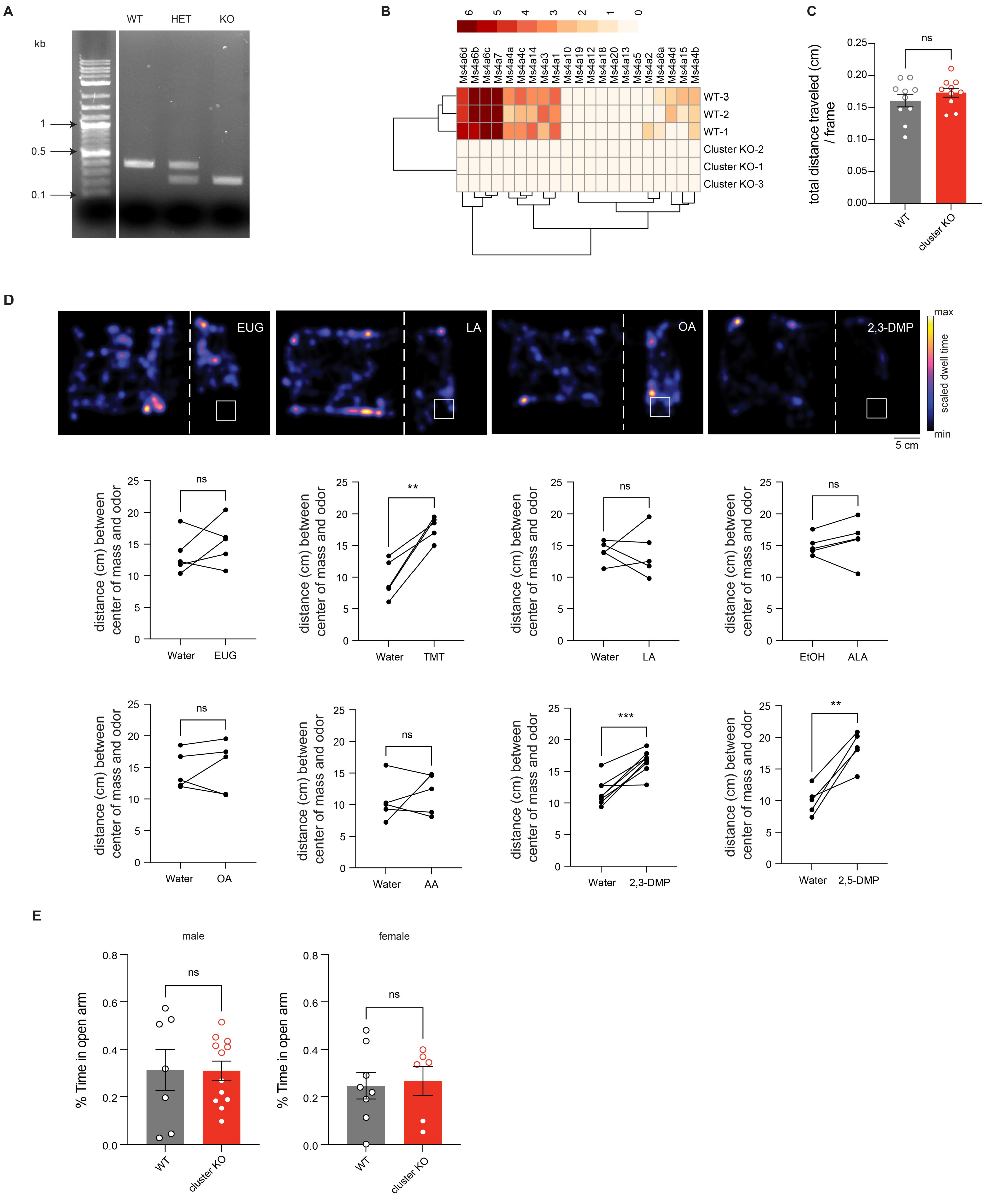

**Figure S2 related to Figure 2**

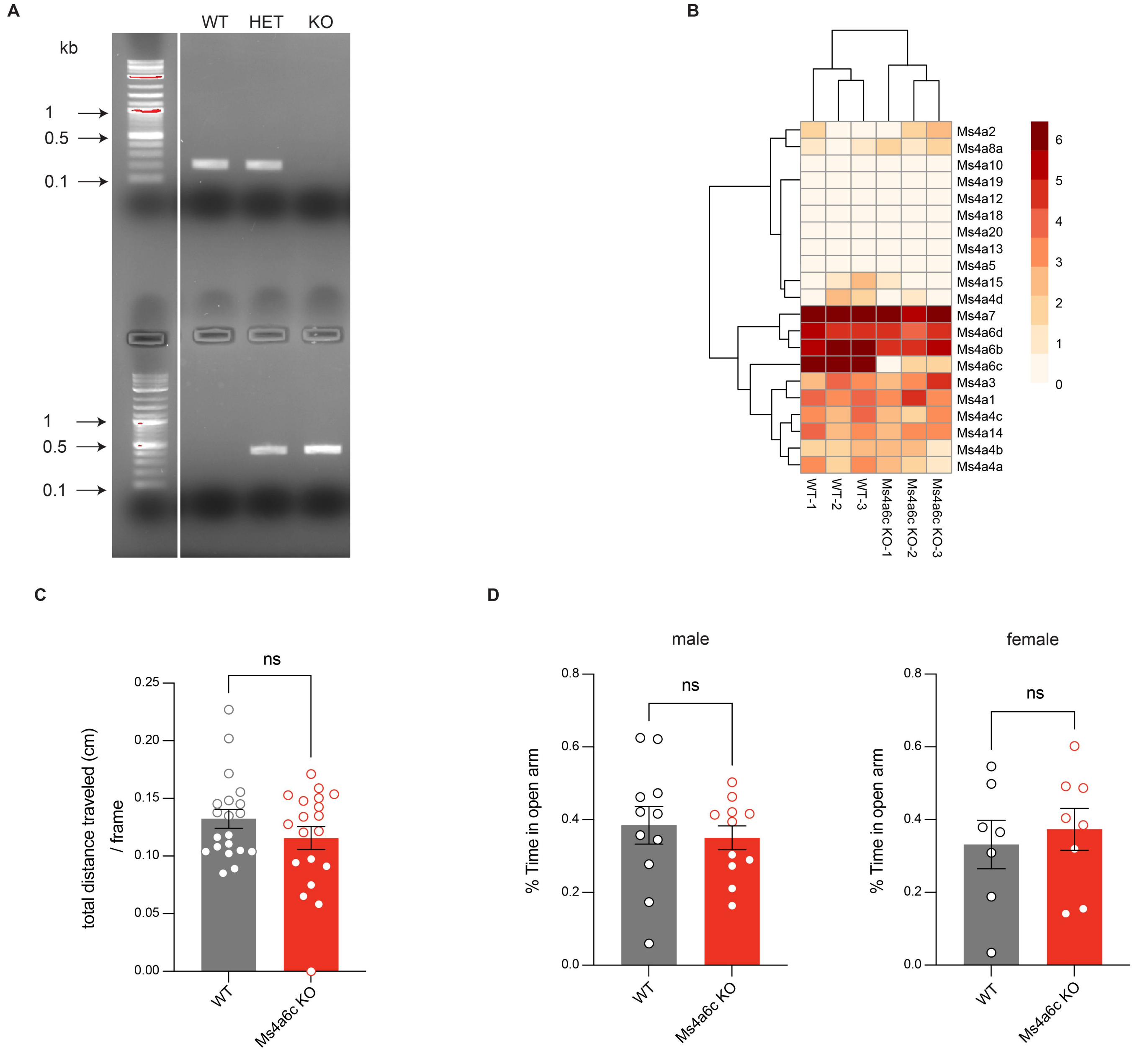

**Figure S3 related to Figure 3**

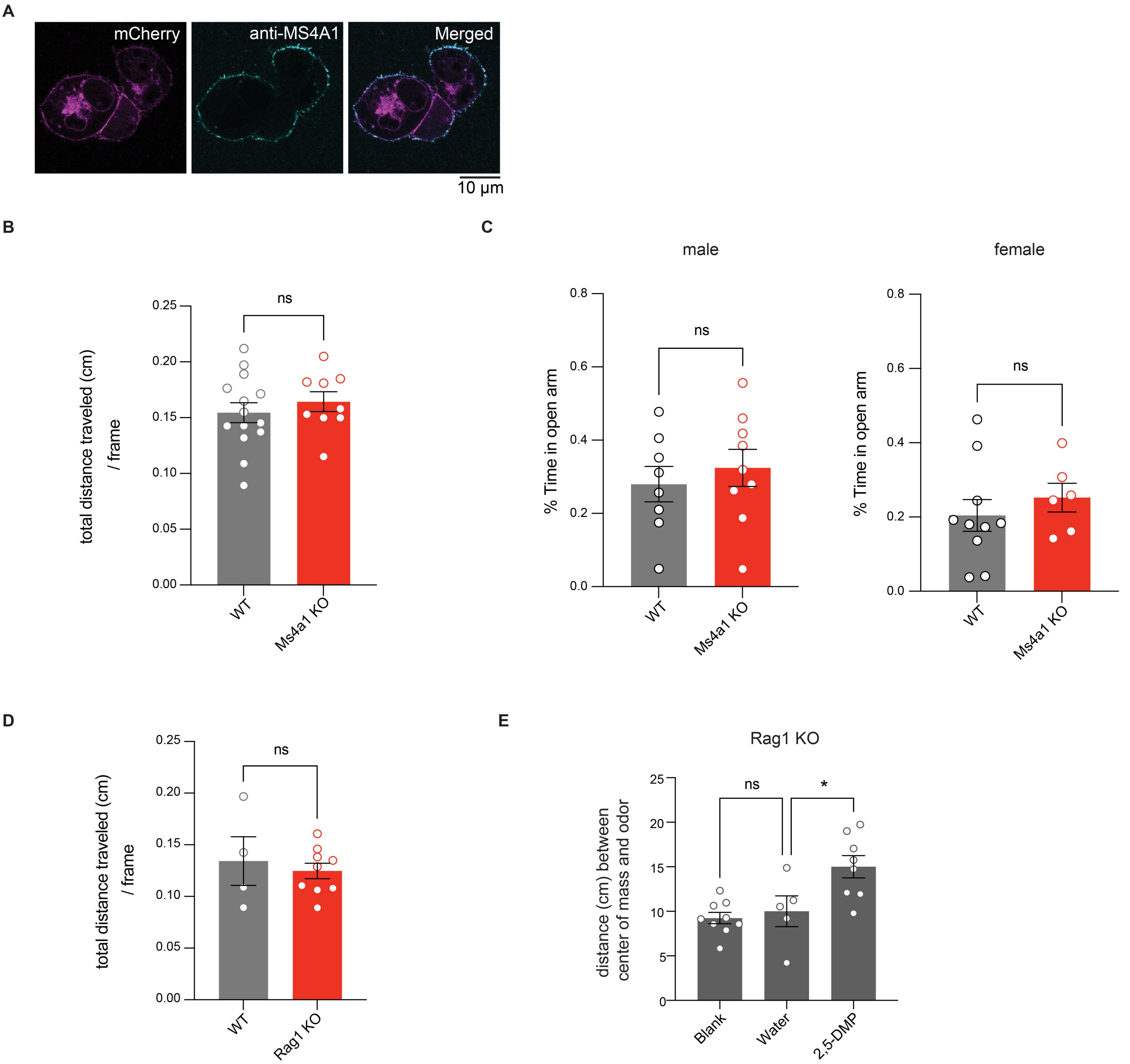

**Figure S5 related to Figure 5**

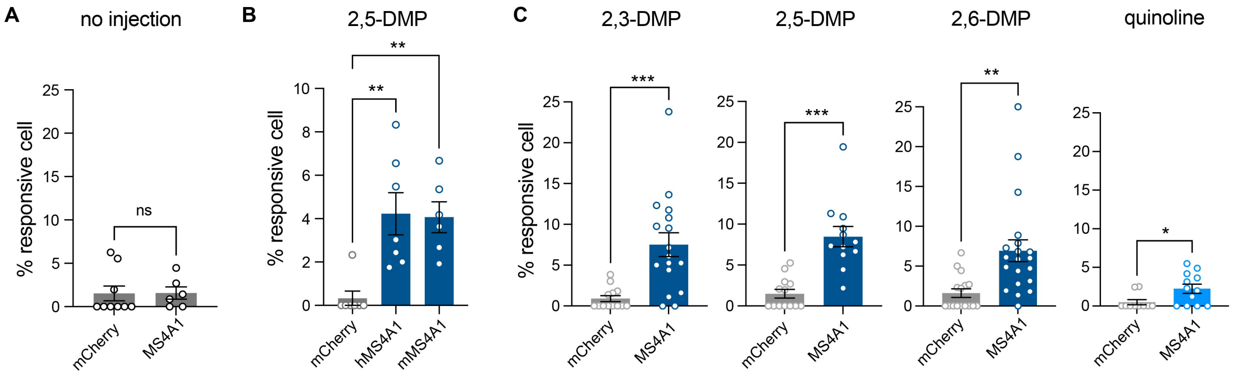
